# A genome-wide CRISPR interference screen using an engineered trafficking biosensor reveals a role for RME-8 in opioid receptor regulation

**DOI:** 10.1101/2022.10.11.511646

**Authors:** Brandon Novy, Hayden Adoff, Monica De Maria, Martin Kampmann, Nikoleta G. Tsvetanova, Mark von Zastrow, Braden Lobingier

## Abstract

G protein-coupled receptors (GPCRs) are the largest family of membrane-bound signaling molecules. Activity of these receptors is critically regulated by their trafficking through the endo-lysosomal pathway. Identifying the genes involved in GPCR trafficking is challenging due the complexity of sorting operations and low affinity protein-protein interactions. Here we present a chemical biology fluorescence-based technique to interrogate GPCR trafficking. We show that the engineered enzyme APEX2 is a highly sensitive biosensor for GPCR trafficking to the lysosome, and this trafficking can be monitored through APEX-based activation of fluorogenic substrates such as Amplex UltraRed (AUR). We used this approach to perform a genome-wide CRISPR interference screen focused on the delta type opioid receptor (DOR), a GPCR which modulates anxiety, depression, and pain. The screen identified 492 genes including known- and novel-regulators of DOR expression and trafficking. We demonstrate that one of the novel genes, RME-8, localizes to early endosomes and plays a critical role in regulating DOR trafficking to the lysosome. Together, our data demonstrate that GPCR-APEX2/AUR is a flexible and highly sensitive chemical biology platform for genetic interrogation of receptor trafficking.

## Introduction

G protein-coupled receptors (GPCRs) control or modulate much of human biology including the physiological responses to many hormones and neurotransmitters (Yang et al. 2021). Consequently, regulation of GPCR signaling is a fundamental cellular task. Membrane trafficking is an important GPCR regulatory mechanism and controls both the subcellular localization and expression levels of the receptor (Marchese et al. 2008; Lobingier and von Zastrow 2019). Broadly, GPCRs traffic through two antiparallel pathways joined at the plasma membrane: a secretory pathway and endo-lysosomal pathway. The secretory pathway controls the flow of new receptors to the cell surface while the endo-lysosomal pathway is an agonist-dependent homeostatic feedback loop to control GPCR levels—and thus cellular sensitivity to receptor activating ligands—through selective receptor proteolysis (Dong et al. 2007; Marchese et al. 2008). GPCRs reach the endo-lysosomal pathway via agonist-stimulated endocytosis and then at endosomes are sorted into pathways that terminate with receptor proteolysis in the lysosome or reinsertion in the plasma membrane (von Zastrow and Sorkin 2021). Which pathway the receptor is sorted into depends on a combination of sorting motifs in the receptor sequence and dynamic post-translation modifications (PTMs) on the receptor (von Zastrow and Sorkin 2021).

A long-standing challenge has been to identify the cellular proteins and pathways which “read-out” this combination of motifs and PTMs into a sorting decision. This has proven difficult due to overall complexity of endosomal sorting operations as well as the low affinity and transient nature of individual protein-receptor interactions that drive these processes (Cullen and Steinberg 2018). While unbiased discovery-based methods have led to important new insights, many of the genes required for selective trafficking and receptor regulation are still unknown (Tanowitz and von Zastrow 2004; Lobingier et al. 2017; Sokolina et al. 2017; Degrandmaison et al. 2020; Semesta et al. 2021).

Genetics is a powerful approach to interrogate intracellular trafficking and identify new genes within these pathways. Our goal was to develop a chemical biology approach that, in conjugation with functional genomics, would provide a screening platform to identify new genes which regulate GPCR trafficking. We focused our chemical biology development on the engineered peroxidase APEX2 (Martell et al. 2012; Lam et al. 2015). We and others have used APEX2 for proximity labeling with GPCRs, but APEX2 can mediate multiple chemistries including activation of fluorogenic substrates including Amplex UltraRed (AUR) (Lobingier et al. 2017; Paek et al. 2017; Civciristov et al. 2019; Pfeiffer et al. 2021; Polacco et al. 2022; Han et al. 2019). We hypothesized that APEX2 could function as a highly sensitive fluorescence-based biosensor for GPCR trafficking to the lysosome.

We selected the delta opioid receptor (DOR) as the first target for our chemical biology and functional genomics approach. The DOR modulates anxiety and depression and is a therapeutic receptor for the treatment of pain (Chu Sin Chung and Kieffer 2013; Gendron et al. 2016; Quirion et al. 2020). Regulation of the DOR involves highly efficient agonist-dependent trafficking to the lysosome, which makes it an excellent model for our approach. Importantly, DOR trafficking plays a key role in the development of drug tolerance to strong agonists (Pradhan et al. 2010). However, the inherent challenges in identifying genes involved in GPCR trafficking have left gaps in our understanding of how the DOR transits the endo-lysosomal pathway and thus which genes contribute to opioid tolerance. Here we demonstrate that a novel chemical biology sensor for lysosomal trafficking—called GPCR-APEX2/AUR—provides a screening platform for identifying new genes that control DOR function through regulated trafficking.

## Results

### APEX2 as a fluorescence-based sensor for membrane protein trafficking to lysosomes

APEX2 is an engineered peroxidase capable of oxidizing structurally distinct substrates (Martell et al. 2012; Lam et al. 2015). Here we were particularly interested in the ability of APEX2 to activate fluorogenic compounds such as Amplex UltraRed (hereafter called AUR) (Zhou et al. 1997; Dwyer et al. 2014; Martell et al. 2017). In this reaction, APEX2 converts the non-fluorescent AUR to a fluorescent resorufin-like molecule in the presence of hydrogen peroxide. All three components are required for the reaction. If APEX2 is the limiting reagent in the system, the assay can be used to quantitatively report enzyme levels. To turn APEX2 into a reporter for DOR expression and trafficking, we inserted a flexible linker and the APEX2 sequence at the DOR C-terminus because that location is amenable to tagging (Scherrer et al. 2006; Henry et al. 2011; Lobingier et al. 2017). In this construct design, APEX2 faces the cytoplasm (pH~7.4) until involution of the DOR-APEX2 into MVBs (pH~6) and subsequent delivery to the lysosome (pH~5) (**Figure 1A**) (Yamashiro and Maxfield 1987; Mitsui et al. 2011; Shen et al. 2013; Johnson et al. 2016; Ostrowski et al. 2016; Webb et al. 2021). We hypothesized that an APEX2-based reporter for lysosomal trafficking would have two advantages: (1) labile to lysosomal proteases and (2) high sensitivity because of multiple substrate turnover events per enzyme.

**Figure 1:**
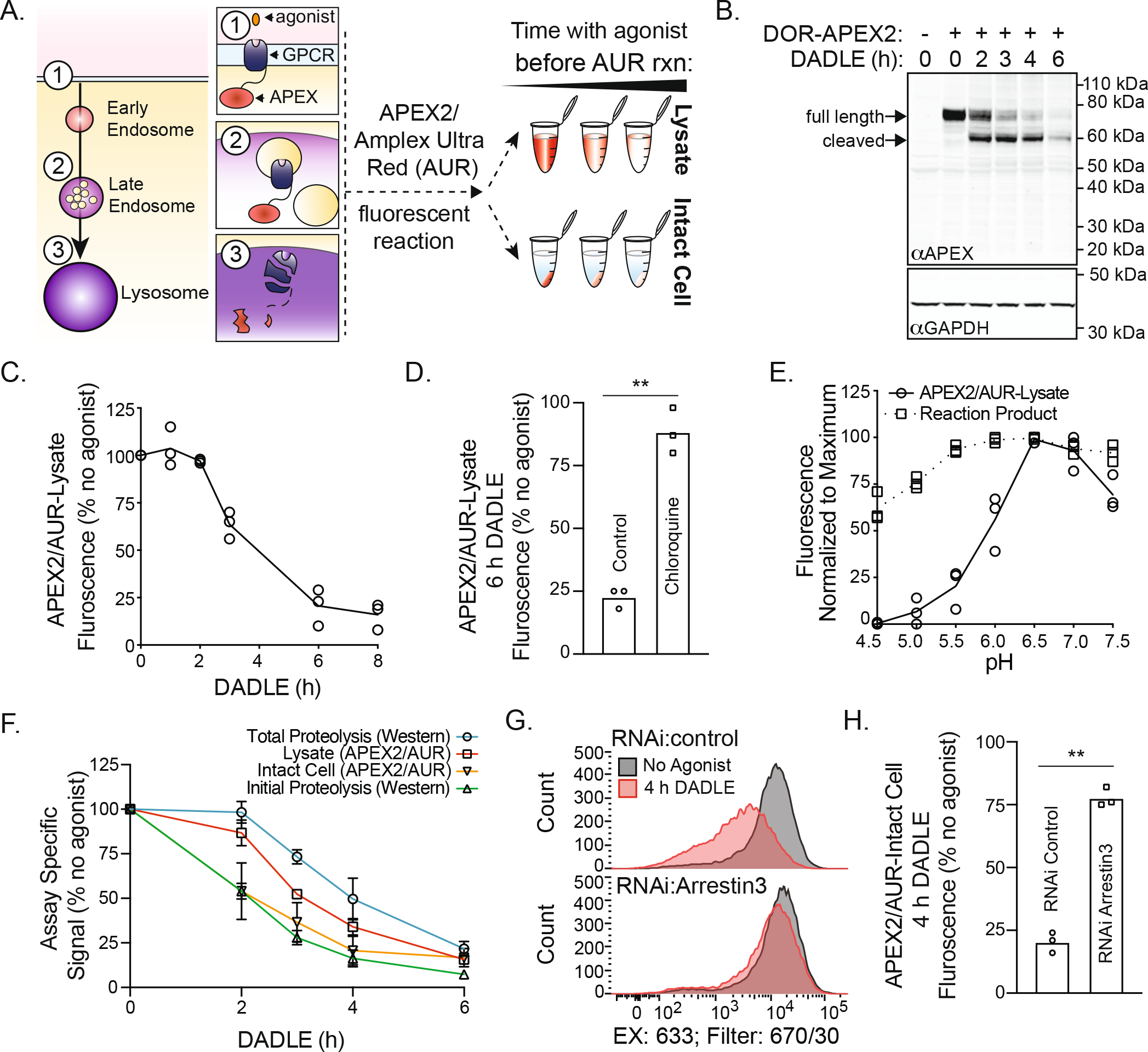
APEX2 as a fluorescence-based sensor for expression and lysosomal trafficking. **(a)** Chemical biology approach using APEX2 and Amplex UltraRed (AUR) as a biosensor for GPCR trafficking to the lysosome. **(b)** Proteolysis of APEX2 detected by western blot following agonist treatment of HEK293 cells stably expressing CMV:DOR-APEX2, n=3, representative example shown, uncropped blot in S1B. **(c)** APEX2/AUR lysate assay in HEK293 cells stably expressing CMV:DOR-APEX2, n=3. **(d)** Chloroquine pretreatment of HEK293 cells stably expressing CMV:DOR-APEX2 followed by agonist stimulation and then the APEX2/AUR lysate assay. N=3 replicates were analyzed with a two-tailed, paired t-test (p=0.0068). **(e)** APEX2/AUR lysate reaction (solid line) performed over pH range in a “universal buffer”. Reaction product (dotted line) denotes data in which the assay was performed at standard pH of 7.4 and then diluted into universal buffer at the noted pH. Data were normalized to the maximum fluorescence (pH 6.5), n=3. **(f)** HEK293 cells stably expressing CMV:DOR-APEX2 were divided and analyzed by western blot or APEX2/AUR assays, n=3. Mean of replicates plotted with SD. **(g, h)** Single cell analysis of APEX2/AUR intact assay following knockdown of Arrestin 3. Fluorescence was measured with a BD FACS Aria II, and a representative example shown. Median fluorescence of single cell populations, n=3, were analyzed with a two-tailed, paired t-test (p=0.0031).

To examine the sensitivity of the approach, we tested its linear range using defined numbers of HEK293 cells stably expressing DOR-APEX2. We found that the APEX2/AUR assay had strong signal from our cell line and a linear range several hundred-fold above background (**Figure S1A**). To test our hypothesis that APEX2 is susceptible to lysosomal proteolysis, we stimulated DOR with its agonist DADLE for different intervals. Probing with an anti-APEX antibody, we found that the DOR-APEX2 transgene (71 kDa) underwent two observable proteolytic events: an initial cleavage of 10 kDa leaving a ~60 kDa product followed by proteolysis of this intermediate and no further observable anti-APEX immunoreactive species (**Figure 1B, S1B**). This time-course is similar to what has been previously observed with a C-terminal tag on the DOR (Hislop et al. 2009; Henry et al. 2011). These data suggest that APEX2, unlike fluorescent proteins, is labile to lysosomal proteolysis (Katayama et al. 2008).

We then tested if APEX2 enzymatic activity could be used as a biosensor for trafficking of the DOR through the endo-lysosomal pathway using the APEX2/AUR assay. To quantify the remaining APEX2 enzymatic activity in cells following treatment with the DOR agonist DADLE, TritonX-100 (0.1% v/v) was used to rapidly permeabilize cellular membranes followed by a 1 minute reaction with AUR and hydrogen peroxide (van de Ven, Adler-Storthz, and Richards-Kortum 2009). The lysate APEX2/AUR reaction was quenched with 100-fold molar excess of a competitive substrate (sodium L-ascorbate) and fluorescence was measured with a plate reader. Consistent with what we observed by western blot, we found that agonist stimulation of cells before the APEX2/AUR reaction decreased formation of the fluorescent product in a time- and agonist-dependent manner (**Figure 1C**). Importantly, pre-incubation with the lysosomotropic compound chloroquine, which increases endosomal/lysosomal pH and decreases activity of lysosomal proteases, strongly blunted the agonist-induced loss of APEX2 activity (**Figure 1D**) (de Duve et al. 1974).

### Comparison of the APEX2/AUR lysate and intact assays

The advantage of the APEX2/AUR assay performed in lysate is that it requires little time (~4 minutes) and minimal material. We thought adapting this method to intact cells would allow single-cell analysis and pooled genetic screens. We developed a version of the APEX2/AUR assay for intact cells using a transient pulse of sodium azide to quench the reaction and bovine serum albumin to scavenge excess dye (Wittrup and Bailey 1988; Burke and Orrenius 1978). For the intact assay, AUR has several advantages: 1) the pKa of its oxidation product is more acidic (~4 vs ~6) than Amplex Red; 2) the non-protonated form of the oxidation product, which we hypothesized would be trapped in cells due to its negative charge, is strongly fluorescent (Ryder, Power, and Glynn 2003; Towne et al. 2004).

We reasoned that one difference between the intact and lysate version of the APEX2/AUR assay would be the pH in the local environment of APEX2. In the intact assay, the endosome and lysosome membranes are not permeabilized with detergent. Thus, during DOR trafficking, the local pH of the APEX2 enzyme would drop following involution of DOR-APEX2 into MVBs and again upon fusion to the lysosome (Yamashiro and Maxfield 1987; Mitsui et al. 2011; Shen et al. 2013; Johnson et al. 2016; Ostrowski et al. 2016; Webb et al. 2021). To directly test APEX2 activity as a function of pH, we performed the lysate assay in a “universal” buffer allowing for a pH range of 4.5-7.5. We measured the optimal pH for APEX2 activity at ~6.9 and a pKa in lysate to be ~5.9 (**Figure 1E**). We also measured the pKa for the fluorescent reaction product of the APEX2/AUR assay and found it to be below 4.5, suggesting it is the activity of APEX2 that is sensitive to mildly acidic conditions (**Figure 1E**).

To test this hypothesis, we directly compared the methods with the same cell population split into the three workflows: western blot, APEX2 activity in lysate, or APEX2 activity within intact cells. We observed that APEX2 activity was lost with t_1/2_ of ~2 hours with the intact assay, and this timing closely mirrored the first proteolytic event observed in western blots (loss of ‘full length’ band in Figure 1B) (**Figure 1F**). Loss of APEX2 enzymatic activity in lysate took longer, with a t_1/2_ of ~3 hours, while the t_1/2_ for loss of all anti-APEX immunoreactivity (both ‘full length’ and ‘cleaved’ bands) was ~4 hours (**Figure 1F**). These results support the hypothesis that, for the intact version of the APEX2/AUR assay, the activity of APEX2 would likely be largely—and quickly—quenched by the pH drop upon involution to MVBs (Yamashiro and Maxfield 1987; Mitsui et al. 2011; Shen et al. 2013; Johnson et al. 2016; Ostrowski et al. 2016; Webb et al. 2021). Comparatively, the permeabilizing conditions of the lysate APEX2/AUR assay appear to reveal a small population of DOR-APEX2 not observed in the intact assay, and we suggest this is likely the fraction of receptor which has undergone involution into MVBs but not undergone significant proteolysis. Together, these data demonstrate that the GPCR-APEX2/AUR approach can be used as a biosensor for agonist-dependent trafficking to the lysosome.

### APEX2/AUR assay is amenable to single-cell analysis using flow cytometry

We then tested the intact cell version of the APEX2/AUR assay using flow cytometry and observed strong fluorescent signal (**Figure S1C**). To test if the intact cell assay was sensitive to agonist-stimulation and on-pathway perturbation, we treated cells stably expressing DOR-APEX2 with siRNAs targeting Arrestin3 (also called B-arrestin2). Arrestin3 is a critical protein for DOR endocytosis, and knockdown of Arrestin3 limits DOR access to the endo-lysosomal pathway (X. Zhang et al. 2008). In cells treated with control siRNA, agonist stimulation before the APEX2/AUR reaction resulted in an ~80% reduction in fluorescence of the cell population while knockdown of Arrestin3 strongly inhibited agonist-induced loss of fluorescence (**Figure 1G, 1H**).

We also tested stability of the fluorescent reaction product within cells using an experimental paradigm in which agonist-treated and unstimulated cells were mixed before the APEX2/AUR reaction. The resulting bi-modal population allows direct comparison of cells with bright or dim signals. We analyzed this cell population by flow cytometry, placed the cells on ice for 30 minutes, and re-analyzed the same material. We found that signal excited by the red laser was stable over that timeframe (APC channel: Ex: 633 nm; Filter: 670/30 nm) (**Figure S1D**). As the oxidation product of AUR has a broad absorbance and can be detected by excitation with multiple laser lines, we also analyzed the behavior of the dye in other channels (**Figure S1D**). We observed less stability of the dye when excited by the 488 nm or 561 nm lasers, and thus it is important with the intact version of the APEX2/AUR assay to use the APC or APC-like channels for analysis and sorting. Together, our work demonstrates that the APEX2/AUR assay can function as a sensitive readout for GPCR expression and lysosomal trafficking and can be utilized in a high-throughput lysate format or for single-cell techniques with intact cells.

### Genome-wide CRISPR interference screen of DOR expression and trafficking

Adapting the APEX2/AUR assay for single-cell work opened the door to pooled genetic screens that use fluorescence activated cell sorting (FACS) to select for phenotype. For our genome-wide screen, we used CRISPR interference (CRISPRi) and second generation libraries encompassing the entire human genome (Qi et al. 2013; Gilbert et al. 2014; Horlbeck et al. 2016). CRISPRi uses a catalytically dead Cas9 (dCas9) fused to the transcriptional repressor KRAB to knockdown gene transcription. CRISPRi has similar efficacy in genome-wide screens to traditional CRISPR-Cas9 with the potential for improved targeting of essential genes (Kampmann 2018; Nagy and Kampmann 2017; Rosenbluh et al. 2017). We created a HEK293 cell line stably expressing both dCas9-KRAB as well as DOR driven by the ubiquitin C promoter (UBC:DOR-APEX2). The UBC promoter expressed 5-fold lower than CMV and, as a human promoter, would be targeted in our genome-wide screen and thus serve as an internal control (**Figure S2A**) (Qin et al. 2010).

We carried out a CRISPRi screen with full coverage of the human genome by independently examining seven sub-libraries consisting of 5 sgRNAs per target gene and a large number of non-targeting controls (NTC) (Horlbeck et al. 2016). We reasoned this would allow us to identify genes with roles in DOR trafficking as well as factors that regulate DOR expression. With the goal of identifying both positive and negative regulators of DOR trafficking, we stimulated cells with agonist for 90 minutes, which was an effective t_1/2_ for the intact APEX2/AUR assay in this line. Following the APEX2/AUR assay, cells were sorted to isolate the top (high, H) and bottom (low, L) quartiles of fluorescence with an estimated sub-library coverage of 500-fold.

Next generation sequencing of the sgRNA-encoding locus was performed to determine the frequency distribution of sgRNAs between the two populations (Kampmann, Bassik, and Weissman 2013). We detected sgRNAs for every gene and the majority had 5 of 5 detected sgRNAs (96.3%), suggesting no major bottlenecks in the workflow (**Table S1**). Using the relative frequency data of the sgRNAs between the two populations, we calculated a knockdown phenotype score (called fold-enrichment: log_2_ (H/L) populations) and significance p-value score for each gene (**Figure 2A, Table S1**). Using the large number of NTCs, we estimated a false discovery rate (FDR) and defined genes with an FDR less than 0.05 as a “hit” (Tian et al. 2019). There were 183 hits with fold-enrichment < 0 (enriched in bottom quartile of fluorescent cells signifying positive regulators of DOR expression or trafficking) and 309 hits with a fold-enrichment > 0 (enriched in top quartile of fluorescent cells signifying negative regulators of DOR expression or trafficking) (**Figure 2A, Table S2**).

**Figure 2:**
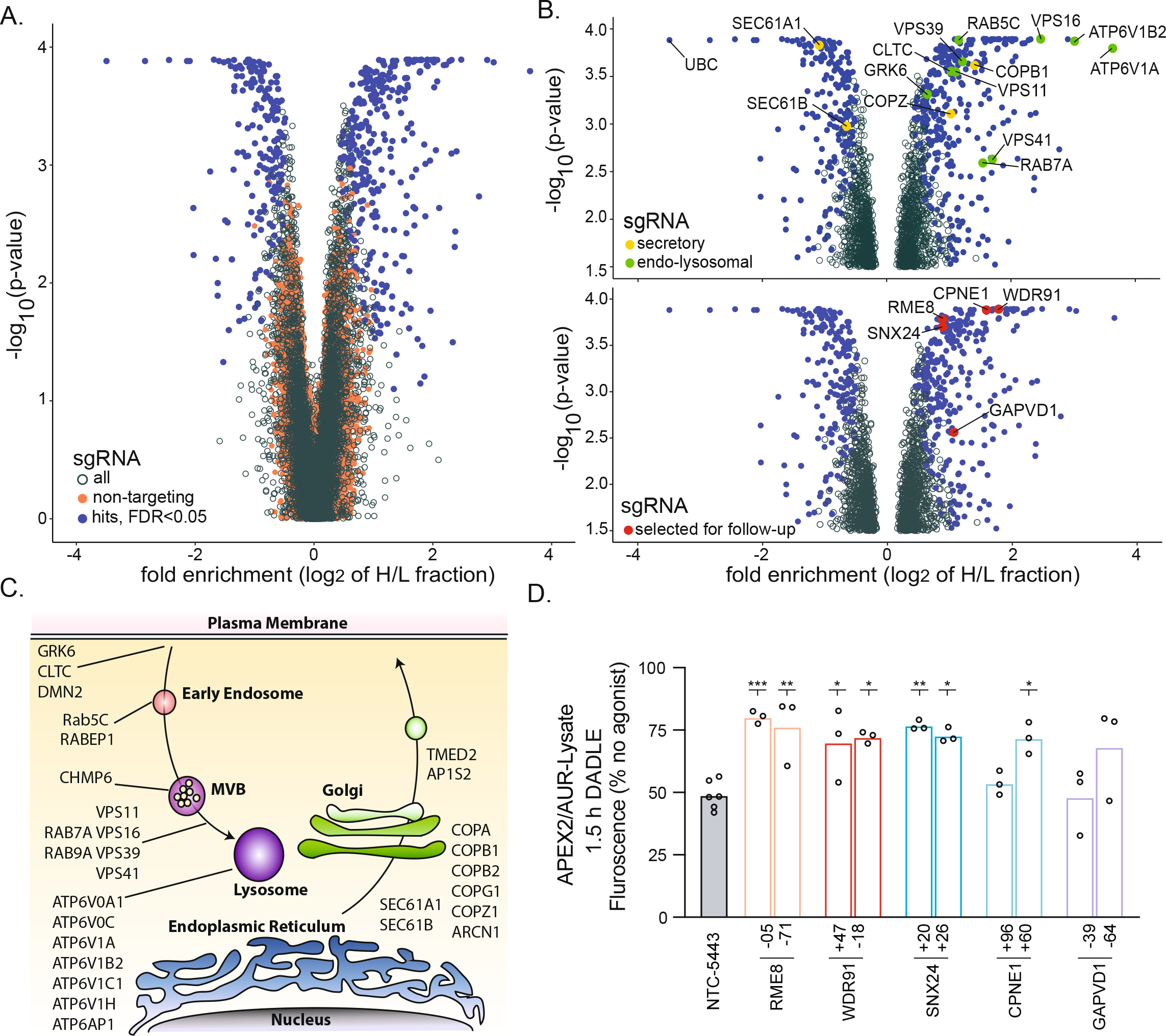
Genome-wide CRISPRi screen for genes affecting DOR expression and trafficking. **(a)** Volcano plot of relative gene enrichment in the high (H) and low (L) signal sorted populations from all seven sub-libraries between FACS sorted bottom and top quartiles following APEX2/AUR intact cell reaction. Genes with an FDR <0.05 are denoted as hits (blue), non-targeting control sgRNAs (orange), and all other gene-targeting sgRNAs (open circles), n=1. **(b)** A subset of hits known to function in secretory or endo-lysosomal are highlighted (upper) and hits selected for follow-up (lower). **(c)** A more comprehensive list of hits known to function in trafficking are diagramed based on primary known location of function. **(d)** HEK293-FLP cells stably expressing UBC:DOR-APEX2, dCas9-BFP-KRAB, and the noted sgRNA, were analyzed using the APEX2/AUR lysate assay. N=3 replicates for all TSS-targeting sgRNAs and n=6 for the NTC. Data were analyzed using an unpaired one-way ANOVA (alpha: 0.05) with Dunnett’s multiple comparisons test. Adjusted p-values when compared to NTC-5433: RME8-05 (0.0008), RME8-71 (0.0032), WDR96+47 (0.0329), WDR96-18 (0.0153), SNX24+20 (0.0027), SNX24+26 (0.0123), CPNE+60 (0.0181).

### Analysis and follow up of the genome-wide CRISPRi screen

We first examined the hits for genes known to function in trafficking. We noted genes linked to endocytic or endo-lysosomal pathways including: endocytic pathway components, endo-lysosomal Rabs, the endosome/lysosome tethering complexes CORVET and HOPS, and subunits of the vacuolar ATPase (**Figure 2B** upper panel and **Figure 2C**). We also identified components of the secretory pathway including subunits of the translocon and coatamer complexes (**Figure 2B** upper panel and **Figure 2C**). As a point of internal validation, sgRNAs targeting the TSS of the ubiquitin C (UBC) gene—which is also the promoter driving DOR-APEX2—were the strongest hit among all sgRNAs enriched low fluorescence population (**Figure 2B**). Thus, the GPCR-APEX2/AUR approach was able to interrogate trafficking at multiple steps of the DOR lifecycle.

To gain a broader perspective on the hits from the screen, we used gene ontology (GO) analysis for cellular components. Consistent with the hits discussed above, we found expected categories related to trafficking including late endosome (GO:0005777) and COPI vesicle coat (GO:0030126) (**Figure S2B**). We also found categories of genes likely to control—directly or indirectly— steady-state expression of the DOR-APEX2 transgene including ribonucleoprotein complex (GO:1990904), replication fork (GO:0005657), and proteasome complex (GO:0000502). Together, these data suggest that the genome-wide screen cast a wide net and identified genes involved DOR expression as well as trafficking.

We then picked five hits for follow-up studies that were not previously known know to function with the DOR but had been described as linked to trafficking: CPNE1, GAPVD1, SNX24, RME-8, and WDR91 (**Figure 2B**, lower panel) (Damer et al. 2005; P. Liu et al. 2018; Hermle et al. 2018; Guillen et al. 2020; Lacey et al. 2022; Fujibayashi et al. 2008; K. Liu et al. 2017; Xing et al. 2021). Two sgRNAs per gene were independently transduced into HEK293-FLP cells stably expressing UBC:DOR-APEX2 and dCas9-KRAB and compared to a NTC sgRNA in agonist-stimulated trafficking of the DOR. We observed that sgRNAs targeting RME-8, WDR91, and SNX24 all inhibited DOR trafficking to the lysosome, with RME-8 knockdown having the largest effects of all sgRNAs examined (**Figure 2D**). We also observed effects with one of the two sgRNAs targeting CPNE1 (CPNE1+60) (**Figure 2D**). Neither of the sgRNAs targeting GAPVD1 had significant effects, although GAPVD1-64 showed a non-significant but apparent trend.

In the process of following up on our genome-wide screen, we noticed that one gene we expected to find—ARRB2/B-arrestin2/Arrestin3—was not a hit with our FDR cutoff of <0.05 (**Table S2**). To follow up on this observation, we selected a single sgRNA from the screen libraries which targets the ARRB2 TSS for independent testing with CRISPRi (Horlbeck et al. 2016). Using the APEX2/AUR lysate assay, we found a small but significant effect when this guide was transduced into the UBC:DOR-APEX2 cell line compared to a NTC sgRNA (**Figure S2C**). This result is consistent with the genome-wide screen data in which ARRB2 showed a moderate phenotypic effect that, when analyzed, did not quite pass the 0.05 FDR threshold (**Table S1**). Together, our follow-up work supported the genome-wide screen, validated three new genes involved in DOR trafficking to the lysosome, and suggested that additional true positive hits exist in the dataset below the selected FDR cutoff. Thus, the GPCR-APEX2/AUR method can be used with CRISPRi to interrogate the human genome and identify novel genes involved in the lifecycle of the DOR.

### RME-8 regulates trafficking of the DOR to the lysosome

One of the strongest hits from our screen and follow up studies was the gene RME-8 (receptor mediated endocytosis 8, also called DNAJC13) (Y. Zhang, Grant, and Hirsh 2001; Girard et al. 2005). RME-8, a gene not found in yeast but conserved from worms to humans, localizes to EEA1-positive endosomes (Y. Zhang, Grant, and Hirsh 2001; Girard et al. 2005; Fujibayashi et al. 2008). RME-8 is necessary for the proper trafficking of recycling cargos and has been shown to physically associate with components of the endosomal recycling machinery including SNX1 and FAM21 (Girard et al. 2005; Shi et al. 2009; Popoff et al. 2009; Freeman, Hesketh, and Seaman 2014). Thus, we were curious why knockdown of RME-8 would have an effect on DOR trafficking to the lysosome. Evidence also demonstrates that RME-8 can regulate the clathrin/HRS/ESCRT degradative subdomain at endosomes via its DnaJ domain and physical interaction with HSC70 (Girard et al. 2005; Shi et al. 2009; Gomez-Lamarca et al. 2015; Xhabija and Vacratsis 2015; Norris et al. 2017; Ryu et al. 2020). Interestingly, the literature is conflicted on whether RME-8 promotes or inhibits trafficking of membrane proteins to the lysosome with studies showing contradictory effects when examining EGFR (Girard and McPherson 2008; Fujibayashi et al. 2008). Intrigued by the links between RME-8 and the recycling and degrading pathways out of the endosome, we sought to further understand the role RME-8 plays in DOR downregulation.

We first asked if RME-8 localizes to DOR-positive endosomes. We identified an effective and specific antibody targeting endogenous RME-8 (**Figure S3A**). Using this antibody, we found that RME-8 puncta were consistently observed as directly adjacent—but with little signal overlap—to DOR positive endosomes (**Figures 3A, 3B**). As a control, we performed additional fixed cell imaging with RME-8 and a component of the recycling pathway, VPS35. We observed a clear colocalization of RME-8 and VPS35 (**Figure 3C, Figure S3B**). We quantified the relative differences with Pearson’s correlation coefficient (PCC) and found higher PCC between RME-8 and VPS35 (0.7±0.1) compared to RME-8 and DOR (0.4±0.1) (**Figure 3D**). To quantify the consistently observed adjacency of RME-8 and DOR signals, we manually scored images for adjacency when <50% RME-8 signal overlaps with proximal DOR or VPS35 signals (see **Figure S3C** for an example of the scoring). We found that 76% of the 518 RME-8/DOR structures quantified showed adjacency while the majority of RME-8/VPS35 signals overlapped and only 35% of the 426 RME-8/VPS35 structures showed adjacency (**Figure 3E**). These data are consistent with the known physical interactions of RME-8 with SNX1 and FAM21 and suggest that in human cells RME-8 is enriched at sites of endosomal tubulation/recycling.

**Figure 3:**
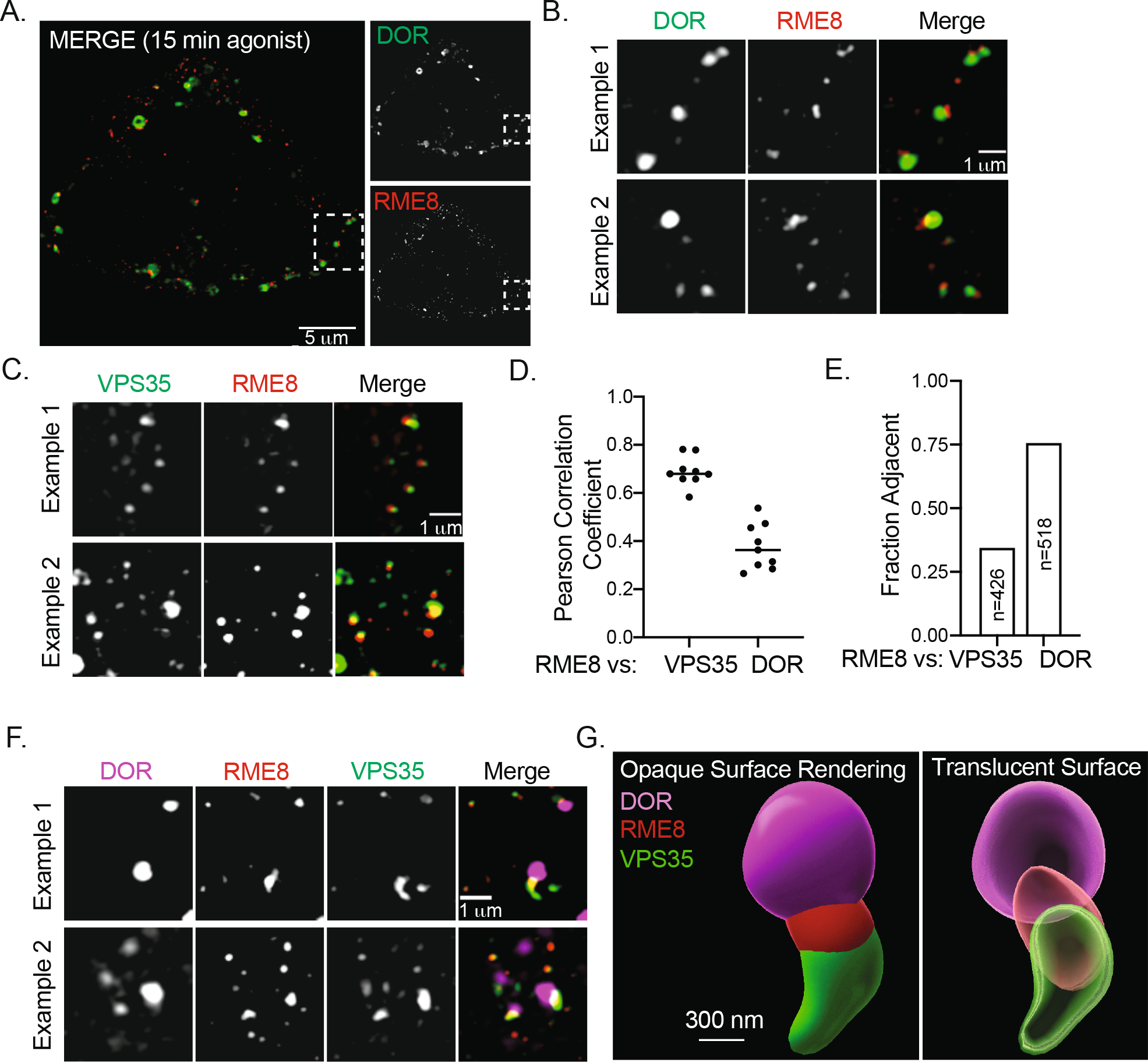
RME-8 localizes to DOR positive endosomes. **(a-c)** Fixed and permeabilized HEK293-FLP cells stably expressing UBC:DOR-APEX2 were probed to detect endogenous RME-8 localization with: (a,b) anti-FLAG (DOR) and anti-RME-8 or (c) anti-VPS35 and anti-RME-8. Scale bar 5 *μ*m in (a) and 1 *μ*m in (b,c). N=3 for all experiments. “Example 1” for 3B comes from white box in 3A while “example 1” for 3C comes from the white box in Figure S3B. In both cases “Example 2” comes from an independent replicate. **(d)** Pearson’s correlation coefficient calculated for DOR/RME-8 or VPS35/RME-8 from n=9 images (3 images per biological replicate). **(e)** Manual scoring of “adjacency” (i.e., <50% signal overlap of the RME-8 with either VPS35 or DOR when both signals appear on the same structure), n=3, individual endosomes counted per condition noted. **(f)** Fixed and permeabilized HEK293-FLP cells stably expressing UBC:DOR-APEX2 were probed to detect endogenous RME-8 with anti-FLAG (DOR), anti-RME-8, and anti-VPS35. Scale bar 1 *μ*m. N=3, “Example 1” comes from white box in S3D, and “Example 2” comes from an independent replicate. **(h)** Imaris software was used to render the surface of a single endosome from the Z-stack images from (3f) “Example 1” and the same model is shown with either an opaque or translucent surface. Scale bar is 300 nm.

To test this hypothesis, we performed three-color imaging experiments with RME-8, DOR to mark the endosome limiting membrane, and VPS35 to mark the recycling domain/recycling tubule. We observed that RME-8 tended to localize to the base of the VPS35-positive recycling tubule on DOR-positive endosomes (**Figure 3F, S3D**). To further illustrate this point, we selected a single endosome with a visually apparent tubule for modeling. Using the information in the z-stack, we created surface renderings of the structure to highlight the relative accumulation of the three proteins (**Figure 3G**). Together, these data suggest that endogenous RME-8 can enrich at sites of endosomal tubulation. Importantly, this observation does not preclude functional interactions between RME-8 and the degradative side of the endosome marked by clathrin/ESCRT domains, and overexpressed RME-8 can localize to entire limiting membrane of the endosome (Girard et al. 2005; Fujibayashi et al. 2008; Shi et al. 2009; Gomez-Lamarca et al. 2015; Xhabija and Vacratsis 2015; Norris et al. 2017).

To more precisely characterize the effects of RME-8 depletion on DOR trafficking to the lysosome, we performed an extended agonist time course. We found that transfecting cells with a pool of four siRNAs targeting RME-8 impeded DOR trafficking to the lysosome, supporting the sgRNAs/CRISPRi results (**Figure 4A**). We then examined each of the four siRNAs making up the pool. We found that the pooled siRNA, as well the individual siRNAs, were able to reduce RME-8 protein levels (**Figure 4B, S4A**). The pooled siRNA and individual siRNAs (#9 and #10) depleted RME-8 protein levels by over 90% and caused significant delay in agonist-induced trafficking of the DOR in the APEX2/AUR assay (**Figure S4B, S4C**). Additionally, we observed no effects from RME-8 knockdown on DOR surface levels, internalization, or recycling (**Figure S4D**), suggesting that the effects of RME-8 knockdown occur while the DOR transits the endo-lysosomal pathway. Of general note, the sensitivity of the APEX2/AUR assay allowed us to observe changes in the baseline rate of DOR degradation in our control conditions. We noted differences in rate between promoters/cell lines (CMV in Figure 1C and UBC in Figure 4A) as well as within a single cell line over time in culture (Figure 4A vs Figure 4D; faster degradation when cells have been in culture longer) (**Figure S4E**). Importantly, the effects of RME-8 knockdown were highly robust and observed in all conditions.

**Figure 4:**
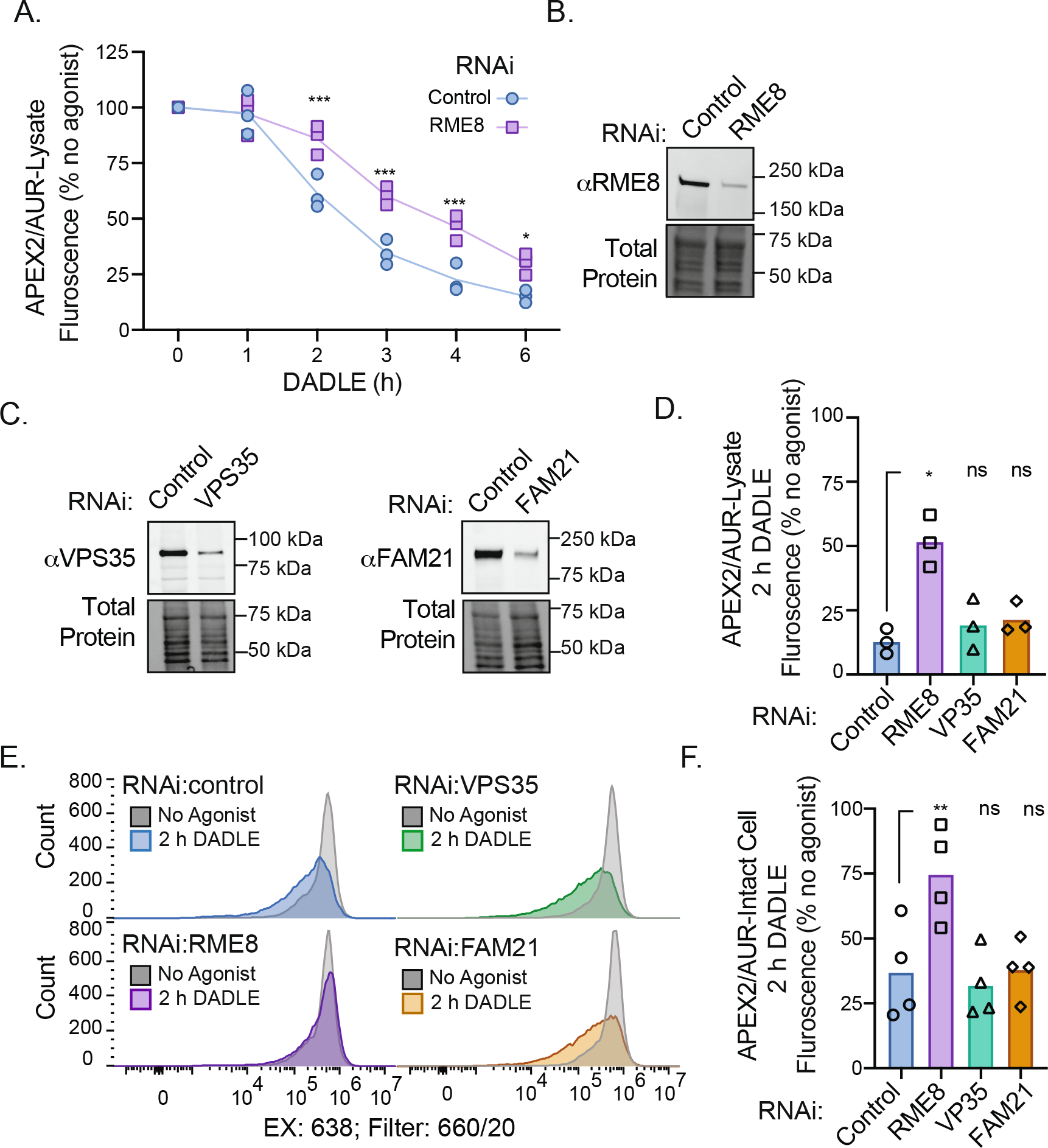
RME-8 regulates trafficking of DOR to the lysosome. **(a)** APEX2/AUR lysate assay following siRNA transfection with HEK293 cells stably expressing UBC:DOR-APEX2. N=3 replicates were analyzed using a multiple t-test with Holm-Sidak correction and an alpha=0.05 (p-values: 2 h=0.0002, 3 h=0.0001, 4 h=0.0002, and 6h=0.0168). **(b,c)** Knockdown of RME-8, VPS35, or FAM21 detected in HEK293-FLP cells stably expressing UBC:DOR-APEX2 by western blot 72 h after siRNA transfection. Total protein stain (Bio-Rad) was used as the loading control, n=3, representative example shown, uncropped blots in S4A and S4E. **(d)** APEX2/AUR lysate assay following siRNA transfection with HEK293 cells stably expressing UBC:DOR-APEX2 with siRNA pool targeting RME-8, VPS35, FAM21A/C, or a non-target control. N=3 replicates were analyzed using a paired one-way ANOVA (alpha: 0.05) with Dunnett’s multiple comparisons test. Adjusted p-values when compared to NTC: RME-8 (0.0352). **(e,f)** Single cell analysis of APEX2/AUR intact assay following knockdown of RME-8, VPS35, or FAM21. The fluorescent reaction product in HEK293 cells stably expressing UBC:DOR-APEX2 was detected with a Beckman Coulter Cytoflex S, representative example shown in (e). N=4 replicates were analyzed using a paired one-way ANOVA (alpha: 0.05) with Dunnett’s multiple comparisons test. Adjusted p-values when compared to NTC: RME-8 (0.0044).

Our data suggest that RME-8 functions to promote DOR trafficking to the lysosome, and we favored the hypothesis that this is due to the known function of RME-8 in regulating the clathrin/ESCRT degradative subdomain through its DnaJ domain and HSC70 (Girard et al. 2005; Shi et al. 2009; Gomez-Lamarca et al. 2015; Xhabija and Vacratsis 2015; Norris et al. 2017; Ryu et al. 2020). However, we wanted to exclude the alternative possibility that perturbation to endosomal recycling following RME8 knockdown could indirectly affect DOR trafficking. To parse which RME-8 modulated pathway contributes to DOR trafficking to the lysosome, we depleted two components of the recycling pathway, VPS35 or FAM21A/C, with siRNA (**Figure S4E**). We observed these pooled siRNAs reduced the levels of VP35 protein by 88% and FAM21 protein by 79% (**Figure 4C, S4F, S4G**). Using the lysate APEX2/AUR assay, we found no significant effects from either VPS35 or FAM21 knockdown on DOR trafficking to the lysosome (**Figure 4D**). To further test this point, we independently performed the same knockdown experiments but analyzed DOR trafficking using the intact APEX2/AUR assay. Again, we observed no significant effects from knockdown of VPS35 or FAM21 on DOR trafficking to the lysosome (**Figure 4E, 4F**). Together, these data suggest that the role of RME-8 in DOR downregulation is due to RME-8 directly modulating the degradative pathway.

## DISCUSSION

Here we describe a novel chemical biology-based reporter for membrane protein expression and trafficking. Our approach leverages the sensitivity of the peroxidase APEX2 and, as a genetic fusion to a GPCR, demonstrates its utility as a biosensor for trafficking to the lysosome. We used this method in combination with a genome-wide CRISPRi screen to identify genes that regulate trafficking of the DOR through its entire lifecycle: from synthesis and insertion at the ER to proteolysis at the lysosome.

### APEX-based chemical biology pipeline to interrogate GPCR function

We have previously used GPCR-APEX2/biotin-phenol to perform sub-minute proximity labeling reactions in living cells (Lobingier et al. 2017). The strength of this approach is its temporal resolution. However, in the context of the GPCR lifecycle, this strength becomes a limitation when trying to control for the multiple organelles through which receptors traffic and the many hours of receptor half-life. The GPCR-APEX2/AUR method presented here addresses this limitation by using the terminal step in the GPCR lifecycle—proteolysis in the lysosome—to integrate the upstream trafficking processes into a single readout. Importantly, the flexibility of the APEX2 enzyme to mediate oxidation of different chemical substrates makes it straightforward to switch between proximity labeling and fluorogenic activation in the same cell line or model system. We anticipate that the addition of AUR approach to the GPCR-APEX2 pipeline will aid in understanding GPCR function at distinct spatiotemporal resolutions.

### Chemical biology reporter for membrane protein trafficking to the lysosome

We found the APEX2/AUR assay has several advantages in analyzing trafficking to the lysosome, particularly compared with gold-standard techniques such as loss of immunoreactivity on a western blot. One advantage of the APEX2/AUR assay is its sensitivity, which arises from the signal amplification created by multiple substrate turnover events per enzyme. While we used stable expression of transgenes in our assays, we speculate that the sensitivity of the APEX2/AUR approach opens the door to inserting the APEX2 tag into an endogenous GPCR locus. A second advantage is the speed and minimal material required to perform the assay in lysate, allowing rapid collection of data and high throughput screening (HTS). A third advantage is the flexibility to perform the assay with intact cells, which allows the use of FACS-based pooled genetic screens. Lastly, while we examined a GPCR that is efficiently targeted for lysosomal proteolysis, we anticipate that our method is amenable to other membrane proteins including those which primarily recycle from endosomes. For such proteins, the APEX2/AUR approach would become a loss-of-signal assay in response to genetic manipulations that block recycling and cause re-routing of the membrane protein to the lysosome (Cao et al. 1999; Temkin et al. 2011; Cullen and Steinberg 2018).

### CRISPR interference screen identifies known and novel trafficking genes regulating the DOR

An advantage of a sensor for the terminal step in the GPCR lifecycle is that it can integrate the secretory, endocytic, and endosomal trafficking steps that precede it. Of the 492 hits we identified, many are genes known to function in membrane protein trafficking including multi-protein complexes such as the secretory COPI complex (6 of 7 subunits identified) or the endo-lysosomal HOPS complex (4 of 6 subunits identified). Thus, our genetic screen likely represents at least a partial “cradle-to-grave” view of the DOR lifecycle. In terms of known players, we were particularly intrigued to find GRK6 as a hit in the screen. GRK6 is a member of GPCR kinase (GRK) family, but previous work has suggested that GRK2/3 are the kinases primarily responsible for DOR endocytosis (Kouhen et al. 2000; Mann et al. 2020; Guo et al. 2000). Additional investigation will be important in defining a role for GRK6 in DOR downregulation.

The driving motivation of our screen was to identify new genes regulating DOR trafficking through the endo-lysosomal pathway. With our primary screen and follow-up studies, we were able to identify three new genes involved in agonist-dependent trafficking of the DOR to the lysosome: RME-8, WDR91, and SNX24. WDR91 is known to function as both a Rab7 effector and regulator of endosomal PI(3)P levels, and knockout of WDR91 inhibits EGFR trafficking to the lysosome (K. Liu et al. 2016, 2017). These known functions are consistent with our observation that loss of WDR91 inhibits agonist-induced trafficking of the DOR to the lysosome. Less is known about the functions of SNX24; like other sorting nexins, SNX24 has a PX domain, and it has also been implicated in alpha-granule formation and MVB biogenesis (Cullen 2008; Lacey et al. 2022). Further studies will be required to define roles for these genes in DOR trafficking.

One intriguing question is how many hits in our screen are specifically involved in the endo-lysosomal trafficking of the DOR? It is a significant experimental challenge to examine hundreds of hits in a standard gene-by-gene format. Additionally, we know from our examination of ARRB2/Arrestin3 that our selected FDR<0.05 excluded true positive hits, and thus it is probable additional candidate genes exist in our data beyond to the 492 highlighted here. One approach that could address this challenge is to utilize small, custom libraries consisting of hundreds of sgRNAs to be analyzed in a pooled format (Nagy and Kampmann 2017; Becker et al. 2020). There are several advantages to such an approach: 1) a reasonably compact micro-library of hits could be analyzed with high coverage (>1000x), allowing greater sensitivity to small effects; 2) the reduced number of sgRNAs compared to the genome-level would allow multiple conditions to be tested in parallel in a reasonable time frame.

### Role of RME-8 in endosomal sorting and DOR trafficking to the lysosome

RME-8 is an intriguing peripheral endosomal protein that has been linked to molecules which regulate both sorting pathways out of the endosome: degradation and recycling. Physically, RME-8 has been shown to interact with components of the recycling pathway including SNX1 and FAM21 (Shi et al. 2009; Freeman, Hesketh, and Seaman 2014). Relatedly, RME-8 depletion has been shown to result in stable, highly branched endosomal recycling tubules, and loss of RME-8 affects the recycling of membrane proteins including CIMPR, STxB, Notch, and MIG-14 (Popoff et al. 2009; Freeman, Hesketh, and Seaman 2014; Gomez-Lamarca et al. 2015; Norris et al. 2017). Yet, RME-8 has also been functionally linked—via its DnaJ domain—with degradation pathway via regulation of clathrin/ESCRT-0 subdomain (Shi et al. 2009; Popoff et al. 2009; Gomez-Lamarca et al. 2015; Xhabija and Vacratsis 2015; Norris et al. 2017). It was not clear how this functional interaction affects membrane protein trafficking to the lysosome, and studies of EGFR trafficking have come to opposite conclusions in terms of a role of RME-8 in sorting of cargo for degradation (Girard and McPherson 2008; Fujibayashi et al. 2008).

Our imaging data demonstrate the majority of endogenous RME-8 localizes near VPS35-positive structures in human cells. The enrichment of RME-8 at the base of recycling tubules is reminiscent of endosomal actin accumulation (Puthenveedu et al. 2010; Shi et al. 2009; Freeman, Hesketh, and Seaman 2014). It is worth noting that while we observed the majority of RME-8 co-localizing with markers of the “recycling domain,” this local enrichment does not preclude RME-8 functioning at other endosomal subdomains. Toward that point, RME-8 has been shown to contain a non-canonical PI(3)P binding domain, which may provide a secondary mechanism for endosomal association and allow it to access the degradative domain (Xhabija and Vacratsis 2015).

Our findings demonstrate that RME-8 is required for efficient sorting of the DOR to the lysosome, and this unlikely to be due to RME-8 function in the recycling pathway. Thus, our data suggest that RME-8 functions to promote degradative sorting at the endosome. The model of RME-8 as a dual-function endosomal protein—regulating proteins mediating recycling or degradative sorting—is consistent with previous observations. Endosomal signaling of the Notch receptor could be perturbed by genetic manipulation of RME-8 in conjunction with either VPS26 (recycling) or HRS (degradation) (Gomez-Lamarca et al. 2015). In *C. elegans*, perturbation of RME-8 resulted in a mixing of degradative subdomain (HRS) and recycling subdomain (SNX1) (Norris et al. 2017). It has previously been proposed that RME-8 recruits HSC70 to endosomes via its DnaJ domain to disassemble endosomal clathrin and thus destabilize HRS/ESCRT-0 (Y. Zhang, Grant, and Hirsh 2001; Shi et al. 2009; Popoff et al. 2009; Gomez-Lamarca et al. 2015; Xhabija and Vacratsis 2015; Norris et al. 2017). If this model is correct, our data would suggest that the outcome of this “pruning” is to maintain a productive degrative endosomal subdomain. What, then, could be the purpose of a dual-function protein such as RME-8? One possibility is that RME-8 functions as a master regulator of endosomal sorting. In such a model, overall cargo flux through the endosome could be accelerated or slowed in response to different cellular states through regulation of RME-8 activity. The exact mechanisms by which this could occur remain to be elucidated.

## Acknowledgements

We thank members of the von Zastrow and Lobingier labs for helpful advice and feedback on this manuscript. We also thank R. Sarah Elmes (UCSF, CA) for critical advice and assistance with flow cytometry.

## Author Contributions

BTL conceptualization; BTL data curation; BTL formal analysis; BTL and MvZ funding acquisition; BN, BTL, HA, MdM, investigation and methodology; BTL, MvZ, NGT, MK resources; MvZ and BTL supervision; BTL writing original draft; BN, MdM, HA, MK, NGT, MzV, BTL review and editing.

## Funding and Additional information

This work was supported by the National Institute of Health (GM137825 and DA043607 to BL, MH109633 to NGT, DA010711 and DA012864 to MvZ, AG062359 to MK). This work was carried out with help of core facility resources: the OHSU Flow Cytometry Core (Matthew Schleisman and Pamela Canaday), the OHSU Advanced Light Microscopy Core (Brian Jenkins and Stefanie Kaech Petrie), the UCSF Center for Advanced Technology (E. Chow), and the UCSF Helen Diller Family Comprehensive Cancer Center Laboratory for Cell Analysis (R. Sarah Elmes; supported by the National Institutes of Health under award P30CA082103).

## Contact for Reagent and Resource Sharing

Requests for resources, reagents, or further information about methods should be directed to Lead Contact Braden Lobingier

## Materials and Methods

### Chemicals

DADLE ([D-Ala2, D-Leu5]-Enkephalin acetate salt) was purchased from Sigma Aldrich (Cat #E7131), resuspend at 10 mM in ddH_2_O, and stored as frozen aliquots at −20 °C. Naloxone hydrochloride was purchased from Tocris (Cat #0599), dissolved in ddH_2_O at 10 mM, aliquoted and stored at −20 °C. Chloroquine diphosphate salt was purchased from Sigma Aldrich (Cat #C6628) and dissolved in ddH_2_O at 100 mM. This chloroquine working stock was filtered, kept at 4 °C for 1 week. Amplex UltraRed was purchased from ThermoFisher Scientific (Cat #A36006), dissolved in anhydrous DMSO with a concentration of 10 mM, aliquoted, protected from light, and stored at −20 °C. Hydrogen peroxide (30% w/w) was purchased from Sigma Aldrich (Cat #H1009) and diluted into ddH_2_O immediately before use. Bovine serum albumin, fatty acid free, was purchased from Sigma Aldrich (Cat #A7030), dissolved in PBS, and filtered before use.

### Antibodies

M1 anti-FLAG was purchased from Sigma Aldrich (Cat #F3040). M1 conjugated to Alexa Fluor 647 (M1-647) with an amine-reactive labeling kit from ThermoFisher Scientific (Cat #A20173). Anti-APEX (anti-L-ascorbate peroxidase 2) was purchased from Abcam (Cat #ab222414). Anti-GAPDH was purchased from EMD Millipore (Cat #MAB374). Anti-RME-8 (also called DNAJC13) was purchased from Fisher Scientific (Cat #70-277-3). Anti-VPS35 was purchased from Novus Biologicals (Cat #NB100-1397).

### siRNA

siRNA targeting ARRB2 (ARRB2-12, Cat #SI03099103) and a non-targeting control (AllStars, Cat # 1027415) was purchased from Qiagen. A pool of 4 siRNAs targeting human RME-8/DNAJC13 was purchased from Dharmacon/Horizon Discovery (Cat #L-010651-01) as well as the individual siRNAs making up that pool: #9 (Cat #L-010651-09), #10 (Cat #L-010651-10), #11 (Cat #L-010651-11), and #12 (Cat #L-010651-12). A pool of 4 non-targeting siRNAs was also purchased from Dharmacon/Horizon Discovery (Cat #D-001810-10). siRNAs were resuspended in either RNase-free water or RNase-free water (Cat# B-003000-WB) supplemented with siRNA buffer (Cat # B-0020000-UB) purchased from Dharmacon/Horizon Discovery.

### cDNA constructs

CMV:DOR-APEX2 is encoded by the construct pcDNA3.1-SSF-DORwt-APEX2 and was previously described (Lobingier et al. 2017). UBC:DOR-APEX2 is encoded by the construct puDNA5-SSF-DORwt-APEX2. puDNA5 was created by using the MluI and ApaI restriction sites to remove the CMV promoter from pcDNA5/FRT and replace it with the human ubiquitin C promoter (UBC). Murine DOR was PCR amplified from pcDNA3.1-SSF-DORwt-APEX2. DOR was cloned into puDNA5 and in-frame with a gBlock encoding a linker (GGGGSGGGG) and APEX2 using In-Fusion to create the final construct, puDNA5-SSF-DORwt-APEX2. dCAS9 is encoded by the construct pHR-SFFV-dCas9-BFP-KRAB and was previously described (Gilbert et al. 2013). sgRNAs were cloned into pU6-sgRNA-EF1alpha-puro-T2A-BFP using the BstXI and BlpI restriction sites (see **Table S3** for a full list of sgRNA sequences and oligos) (Gilbert et al. 2014)

### Cell Culture and Stable Cell Line Generation

HEK293 cells, and FLP-In-293 (HEK293-FLP) cells purchased from Thermo Fisher Scientific (Cat # R75007), were cultured in DMEM (Cat #11965-092) supplemented with 10% FBS and grown at 37 °C with 5% CO_2_. HEK293 cells stably expressing CMV:DOR-APEX2 were created as previously described and maintained in 1:1000 G418/Geneticin (Thermo Fisher Scientific, Cat # 10131035) (Lobingier et al. 2017). HEK293 Flp-In cells stable expressing UBC:DOR-APEX2 were created by transient transfection of puDNA5-SSF-DORwt-APEX2 and pOG44 using Lipofectamine 2000 (ThermoFisher Scientific, Cat # 11668019). To ensure sufficient genetic diversity, a ~40% confluent T75 was transfected to ensure >10 ‘plaques’ grew upon addition and selection with 100 *μ*g/mL hygromycin (ThermoFisher Scientific, Cat #10687010). Following transfection and selection, all hygromycin-resistant HEK293 Flp-In cells were sorted for expression, expanded, and frozen for further experimentation. HEK293 Flp-In cells expressing UBC:DOR-APEX2 were maintained in 50 *μ*g/mL hygromycin. To generate HEK293 cell lines stably expressing dCas9, Lenti-X HEK293 cells (Takara Bio, Cat #632180) were transfected with pHR-SFFV-dCas9-BFP-KRAB, pVSVG, and psPAX2 using Lipofectamine 2000. Supernatant was harvested after either 48 or 72 h, filtered through a 0.45 *μ*m PES filter, and incubated overnight with HEK293 FLP-In cells stably expressing UBC:DOR-APEX2. Cells expressing both DOR and dCas9 were isolated using fluorescence activated cell sorting (FACS) on a BD FACS Aria 2 (BD Biosciences), serially diluted, and expanded as individual clones. Clones were validated for expression of DOR, dCas9, and activity of dCas9 using sgRNAs targeting the TSS of ARRB2.

### siRNA transfections

Knockdown of ARRB2/Arrestin3 in CMV:DOR-APEX2 expressing cells was performed using “reverse” transfection in which 20 pmol siRNA (Qiagen) and 12 *μ*L RNAimax (ThermoFisher Scientific, Cat #13778075) were incubated for 20 minutes in OptiMem (ThermoFisher Scientific, Cat # 31985070) and then added to a ~50% confluent cell suspension in a 6 cm plate. Media was exchanged 24 h after transfection, cells were lifted and replated 48 h after transfection, and experiments performed 72 h after transfection.

Knockdown of RME-8 in UBC:DOR-APEX2 expressing cells was performed using “reverse” transfection in which 19.8 pmol siRNA (Dharmacon) and 3.33 *μ*L Dharmafect1 were incubated for 20 minutes in OptiMem and then added to a ~20% confluent cell suspension in one well of a 6-well plate. Cells were lifted and replated 24 h after transfection, and experiments were performed 72 h after transfection. Knockdown of VPS35 and RME8 for Figure 4 (C-F) was performed using “reverse” transfection in which 39.6 pmol siRNA (Dharmacon) and 6.66 *μ*L Dharmafect1 were incubated for 20 minutes in OptiMem and then added to a ~20% confluent cell suspension in one well of a 6-well plate. Cells were lifted and replated 24 h after transfection, and experiments were performed 72 h after transfection. The above protocol was employed for FAM21 utilizing 19.8 pmol of each siRNA pool targeting FAM21 B and C respectively.

### Western Blot

Western blots were performed with slight variations depending on the experiment. For HEK293 cells stably expressing CMV:DOR-APEX2, the cell pellet was directly lysed in RIPA (50 mM Tris, 150 mM NaCl, 1% TritonX-100, 0.5% sodium deoxycholate, 0.1% SDS, pH 7.4) supplemented with a protease inhibitor cocktail (Millipore Sigma, Cat #11836170001), incubated on ice for 10 minutes, sonicated, and centrifuged at 10,000 x g for 10 minutes. Supernatant was mixed with sample loading buffer supplemented with 1% v/v 2-mercaptoethanol (BioRad, Cat #1610710). Sample was not heated to prevent aggregation of the GPCR. Proteins were separated on a NuPAGE 4-12% Bis-Tris Gel (ThermoFisher Scientific, Cat # NP0321) in NuPAGE MOPS buffer (ThermoFisher Scientific, Cat # NP0001). Proteins were transferred to nitrocellulose (BioRad, Cat #162-0112) blocked with Odyssey Blocking Buffer (LICOR, Cat # 927-50000) for 1 h, and probed with primary antibody for 2 h at room temperature. Membranes were washed with TBS supplemented with 0.1% v/v Tween, incubated with 680-donkey-anti-rabbit (LI-COR, Cat #926-68073) or 800-donkey-anti-mouse (LI-COR, Cat #926-32212) at 1:3000 for 1 hr, and then imaged on an Odyssey Infrared Imaging System (LI-COR). For quantification, indicated band intensity was analyzed using ImageStudioLite (LI-COR) with a non-expressing HEK293 control for background subtraction, and normalized to the GAPDH loading control.

For HEK293-FLP cells stably expressing UBC:DOR-APEX2: cells were lifted and spun down and then lysed in RIPA (50 mM Tris, 150 mM NaCl, 1% TritonX-100, 0.5% sodium deoxycholate, 0.1% SDS, pH 7.4) supplemented with a protease inhibitor cocktail, incubated on ice for 10 minutes, sonicated, and centrifuged at 10,000 x g for 10 minutes. Supernatant was mixed with sample loading buffer supplemented with Invitrogen Bolt reducing agent. Sample was not heated to prevent aggregation of the GPCR. Proteins were separated on a 4–20% Mini-PROTEAN TGX Stain-Free™ Protein Gels in SDS-PAGE Running Buffer (0.2501 M Tris, 1.924 M Glycine, 0.0347 M SDS). After performing a 45 second UV-activation for the stain-free total protein stain, proteins were transferred to nitrocellulose blocked with Everyblot Biorad Blocking Buffer for 1 h, and then probed with primary antibody overnight at 4 °C. Membranes were washed with PBS supplemented with 0.1% v/v Tween, incubated with StarBright Blue 700 (or 520) Goat Anti-Rabbit or StarBright Blue 700 (or 520) Goat Anti-Rabbit at 1:2000 for 1 h at room temperature. After washing, the membranes were imaged on a Biorad Chemidoc Imaging System. For quantification, indicated band intensity was analyzed using ImageLab (Biorad) or ImageJ using total protein stain for normalization. siRNA knockdown efficacy was assessed comparing band intensity after normalization to a non-targeting control.

### Amplex Assay, Lysate

The lysate assay was performed with slight variations depending on the experiment. For Figure 1C, 1D, and 1G: HEK293 cells stably expressing CMV:DOR-APEX2 were either unstimulated or stimulated with 10 *μ*M DADLE for indicated duration. For experiments testing the effect of chloroquine, chloroquine was added 30 min prior to agonist stimulation at 100 *μ*M final concentration. Cells were washed 1x with PBS supplemented with EDTA and detached from the 6 well plate with PBS+EDTA. Cells were pelleted with centrifugation, 500 x g for 5 minutes. Reaction buffer #1 (PBS, 0.1% TritonX-100, 100 *μ*M Amplex UltraRed, 5 mM H_2_O_2_ and protease inhibitor cocktail) was added to cell pellet on ice. The reaction was allowed to proceed for 2 minutes at room temperature, and stopped by directly adding 10 mM final of freshly prepared sodium L-ascorbate. For 1C and 1D fluorescence measurements, excitation 485 and emission 590, were taken using a Spectra Max Gemini XPS (Molecular Devices). For 1G fluorescence measurements, excitation 561 nm and emission 610 nm, were taken using a Spectra Max Gemini XPS (Molecular Devices).

For Figures 2D, S2C, 4A, 4D, S4C: HEK293-FLP cells stably expressing UBC:DOR-APEX2 were either unstimulated or stimulated with 10 *μ*M DADLE for the indicated duration. No differences were noted in experimental outcome between these differing conditions. For experiments testing RME-8 knockdown, the cells were plated into a 24-well plate following siRNA treatment as described above. Cells were lysed in well using ice-cold Lysis Buffer (PBS, 0.1% TritonX-100) which was allowed to proceed for 3 minutes at room temperature. Reaction buffer 2x (PBS, 0.1% TritonX-100, 200 *μ*M Amplex UltraRed, 200 *μ*M H_2_O_2_) was then added on top of the Lysis Buffer and allowed to proceed for ~1 minute. The reaction was stopped by adding Quench buffer 3x (PBS, 30 mM sodium L-ascorbate). Fluorescence measurements, excitation 555 nm and emission 610 nm, were taken using a Spark multimode microplate reader (Tecan).

### Linear Range of APEX Assay, Lysate

HEK293-FLP cells stably expressing UBC:DOR-APEX2 were washed with PBS, detached with TrypLE Express (ThermoFisher Scientific, Cat # 12604039), and centrifuged with an equivalent volume of DMEM+10% FBS. Cells were counted in Trypan Blue (ThermoFisher Scientific, Cat #15250061) using a Countess FL (Invitrogen) and concentration was determined as the average counts from two areas on both sides of a hemocytometer. Cells were aliquoted into separate tubes spanning a range of 39,250-1,256,000 cells per reaction. Cell pellets were lysed for 3 min on ice in PBS+0.1% TritonX-100. 2x reaction buffer (PBS, 0.1% TritonX-100, 200 *μ*M Amplex UltraRed, 200 *μ*M H_2_O_2_) was added and the reaction was allowed to proceed for 1 minute before addition of 3x Quench Buffer (PBS, 30 mM sodium L-ascorbate). Quenched lysate was then moved to a 96 well plate (Costar, black half area, Cat # 3694) and fluorescence was measured, excitation 561 +/− 10 nm and emission 610 +/− 10 nm, using a Spark multimode microplate reader (Tecan) with gain optimized each run to the highest signal sample.

### Amplex Assay, Intact Cells

For Figures 1G, 1H, S1C, and S1D, HEK293 cells stably expressing CMV:DOR-APEX2 were washed with PBS supplemented with EDTA, detached from the plate with TrypLE Express, and pelleted with an equivalent volume of DMEM+10%FBS by centrifugation. Cells were resuspended in ice-cold PBS supplemented with 200 *μ*M Amplex UltraRed, incubated at room temperature for 5 minutes followed by 10 minutes on ice. To avoid the potential of any extracellular reaction products interfering with our measurements, the reactions prepared for FACS were performed in the presence of bovine serum albumin (BSA) which has been shown to be highly effective at capturing extracellular resorufin and resorufin-like molecules (Wittrup and Bailey 1988; Burke and Orrenius 1978). 2x Reaction buffer (PBS, 4% w/v BSA, 100 *μ*M H_2_O_2_) was prepared immediately before use and added directly to cells. Reaction was allowed to proceed for 30 seconds and quenched with the addition of sodium azide, 1 mM final. Cells were pelleted with centrifugation at 500 x g for 1 minute, washed with PBS supplemented with 2% BSA, and then resuspended in FACS buffer (PBS supplemented with 1% BSA). Cells were passed through a 40-*μ*m mesh nylon filter (Corning, Cat #352340) prior to analysis on a BD FACS Aria using the following channels: APC (Excitation: 633 nm; Emission:670/30 nm), mCherry (Excitation:561 nm; Emission: 610/20-600 nm LP); PE (Excitation: 488 nm; Em: 575/25-500 nm LP), Pacific Blue (Excitation: 405 nm; Emission: 450/50 nm)). At least 20,000 cells were counted per sample, and singlets were gated for further analysis. For analysis of percent signal loss in RNAi treated cells, median fluorescence of the population was determined using FlowJo and background subtracted from a population in which no APEX2/AUR was performed.

For Figures 4E and 4F, HEK293 cells stably expressing CMV:DOR-APEX2 were treated with siRNA following the siRNA transfection protocol above. To assess DOR-trafficking, cells were either unstimulated or stimulated with 10 *μ*M DADLE for 2 hours and assayed in accordance with the above protocol, without a final filtration step. Analysis was conducted on Cytoflex S (Beckman Coulter) using the following channel: APC (Excitation: 638 nm; Emission:660/20 nm). At least 15,000 cells were counted per sample, and singlets were gated for further analysis. For analysis of fluorescence remaining, the geometric mean of both stimulated and unstimulated populations were obtained for each siRNA using FlowJo.

### APEX activity as a function of pH

The lysate APEX2/AUR assay was performed as described above except that the HEK293 cells stably expressing CMV:DOR-APEX2 were directly lysed with Universal Buffer (20 mM sodium acetate, 20 mM MES, 20 mM HEPES, 100 mM sodium chloride, 0.1% TritonX-100) at the indicated pH for 3 minutes on ice. 2x Reaction Buffer #2 (Universal buffer supplemented with 200 *μ*M Amplex UltraRed, 100 *μ*M H_2_O_2_) was added directly to the lysis and allow to proceed for 1 minute. The reaction was stopped by the addition of 10 mM final sodium L-ascorbate. To measure the pH dependence of the oxidation product, universal buffer was replaced with PBS for the reaction and the quenched reaction product was added as a 1:10 dilution to the Universal Buffer at the indicated pH. Fluorescence measurements, excitation 561 nm and emission 610 nm, were taken using a Spectra Max Gemini XPS (Molecular Devices). Data were processed by subtracting fluorescence from the buffer alone and normalized to the maximal fluorescence observed for the assay, at pH 6.5.

### Comparison of Amplex Assay (Lysate vs Intact) and Western blot

HEK293 cells stably expressing CMV:DOR-APEX2 were either unstimulated or stimulated with 10 *μ*M DADLE for 2, 3, 4, or 6h. Cells were washed 1x with PBS+EDTA, detached from the plate with TrypLE Express, and split into three different workflows: lysate APEX2/AUR, intact cell APEX2/AUR, and western blot assays. Fluorescence of both lysate and intact cell assays measured with a Spectra Max Gemini XPS (Molecular Devices), excitation 561 nm and emission 610 nm. Western blot data was acquired using an Odyssey Infrared Imaging System (LI-COR) and quantified with ImageStudioLite (LI-COR).

### Effect from RME-8 knockdown on Surface expression, Internalization, and Recycling Assays

HEK293-Flp cells stably expressing UBC:DOR-APEX2 were transfected with pooled siRNAs targeting RME-8 or a non-targeting control as described above. 72 h after transfection, cells were stimulated with 10 *μ*M DADLE for 30 minutes (internalization), 10 *μ*M naloxone for 30 minutes (surface), or 30 minutes of 10 *μ*M DADLE, one media wash, followed by 30 minutes of 10 *μ*M naloxone (recycling). Cells were washed 1x with PBS (Corning, Cat #21-031-CV), detached from the plate with TrypLE Express, and resuspended in FACS-Buffer #2: PBS with calcium and magnesium (Corning, Cat #21-030-CV) supplemented with 1% BSA. Alexa647-conjugated M1-anti-FLAG antibody was incubated with cells for 1 h at 4 °C, washed 1x, and resuspended in FACS-Buffer #2. Cells were analyzed in a Cytoflex S (Beckman Coulter) using the following channel: APC (Excitation: 638 nm; Emission:660/20 nm). At least 10,000 cells were counted per sample, and singlets were gated for further analysis. The geometric mean of the APC channel was used to quantify surface expression, internalization (1-[agonist/antagonist]), or recycling ([recycling-agonist]/[antagonist-agonist]).

### Comparison of DOR surface expression between CMV and UBC promoters

HEK293-Flp cells stably expressing UBC:DOR-APEX2 or CMV:DOR-APEX2 were seeded in addition to the parental line and cultured for 24 hours. Cells were approximately 80% confluent when washed 1x with PBS (Corning, Cat #21-031-CV), detached from the plate with TrypLE Express, and resuspended in FACS-Buffer #2: PBS with calcium and magnesium (Corning, Cat #21-030-CV) supplemented with 1% BSA. Alexa647-conjugated M1-anti-FLAG antibody was incubated with cells for 1 h at 4 °C, washed 1x, and resuspended in FACS-Buffer #2. Cells were analyzed in a Cytoflex S (Beckman Coulter) using the following channel: Excitation: 638 nm; Emission:660/20 nm. At least 10,000 cells were counted per sample, and singlets were gated for further analysis. The geometric mean of the APC channel was used to quantify surface expression with the parental line establishing a baseline control to subtract background fluorescence ([UBC – Parental] / [CMV – Parental]).

### Genome-wide CRISPR interference screen

HEK293-FLP cells stably expressing UBC:DOR-APEX2 and dCas9-BFP-KRAB were transduced, separately, with seven different CRISPRi sgRNA sub-libraries to obtain coverage of the entire human genome (Horlbeck et al. 2016). To ensure the majority of transduced cells only contained a single sgRNA (MOI < 1), initial transduction rates were kept below 40% with sufficient cells transduced and grown to ensure >500-fold coverage of the sub-libraries. Two days after transduction, puromycin (Gibco, Cat #A1113803) at a final concentration of 0.75 *μ*g/mL was added for 72 h to select for transduced cells. Cells were allowed to recover and grown for an additional two days. At 6 days post-transduction, cells were stimulated with DADLE for 90 minutes and the intact cell version of the APEX2/AUR assay was performed as follows: following agonist stimulation, cells were washed with PBS+EDTA, detached with TrypLE Express, and centrifuged with an equivalent volume of DMEM+10% FBS. All cells were resuspended in PBS, pooled, divided into 8 fractions, and then processed 4 fractions at a time. 200 *μ*M AUR was added directly to foil-covered cells, incubated for 5 minutes at RT, and then 10 minutes at 4 °C. 2x Reaction buffer #4 (PBS+4% w/v BSA +100 *μ*M H_2_O_2_) was added to cells, reaction was allowed to proceed for 90 seconds, and then quenched through addition of sodium azide (1 mM final). Cells were centrifuged for 3 minutes at 500 x g at 4 °C, washed 1x with PBS +2% BSA, and resuspended in PBS+1% BSA. All samples were pooled, passed through a 0.45 um nylon filter, and prepared for FACS on a BD FACS Aria 2 using the following nested gating scheme: singlets, BFP positive, and the top and bottom quartile of the APC channel. Genomic DNA was harvested from the FACS sorted cells using QIAamp DNA Blood Maxi kit (Qiagen, Cat #51192), sgRNA libraries were prepared using Q5 HotStart High Fidelity (NEB, Cat #M0493L) and barcoding primers (**Table S4**). PCR products were purified using QIAquick PCR purification columns (Qiagen, Cat #28106), loaded onto 20% TBE gels (ThermoFisher Scientific, Cat #EC63155BOX), and a ~270 bp band was excised from the gel. This product was quantified using a Bioanalyzer (Agilent) and sequenced on an Illumina HiSeq-4000 (Illumina) using custom primers (**Table S4**). One biological replicate was performed of this screen.

### Bioinformatic Analysis of CRISPRi screen

Next generation sequencing data from the CRISPRi screen was analyzed as previously described (Kampmann, Bassik, and Weissman 2014; Semesta et al. 2021). Each sublibrary was analyzed independently as follows: raw sequencing reads were cropped and aligned and filtered by expected libraries. A phenotype score, called fold-enrichment (also called epsilon), was calculated as a log2 ratio using the three strongest sgRNAs per TSS by absolute value. P-values were calculated using a Mann-Whitney test comparing all five sgRNAs targeting a given TSS relative to all non-targeting controls. A composite score (the product of the epsilon and p-value) was used to calculate an FDR with a cutoff of <0.05 and identify high-confidence “hits” as described previously (Tian et al. 2019).

### GO code analysis

The 492 high confidence hits from the screen were analyzed using the PANTHER overrepresentation analysis for “Slim Cellular Component” with a Fisher’s exact test and Bonferroni correction for multiple comparisons. GO categories with statistically significant overrepresentation were ranked by the number of hits found, and any redundant category (defined as a category with only one unique hit relative to previous categories) were removed. Overly broad categories (e.g., “cellular_component”) were also removed.

### sgRNA follow up screen

To generate lentivirus for sgRNAs targeting select gene TSS, LentiX HEK293T cells were transfected using Lipofectamine 2000 and the pU6-sgRNA of interest, pPAX2, and pVSVG. Media on 293T cells was changed 24 h after transfection, and viral supernatant was collected and filtered at 48 h after transfection. HEK293-FLP cells stably expressing UBC:DOR-APEX2 and dCas9-BFP-KRAB were transduced, separately, with viruses encoding the different sgRNAs. 2 *μ*g/mL puromycin was added 24 h after transduction to select cells with sgRNAs, and selection was maintained for 3 passages across 7 days. Cells were frozen and thawed to be assayed over ~10 days, after which a new aliquot was thawed. Cells were analyzed using the lysate APEX2/AUR assay as described above.

### Fixed Imaging

HEK293-FLP cells stably expressing UBC:DOR-APEX2 were plated onto poly-L-lysine (Sigma Aldrich, Cat # P8920-100ML) coated coverslips in wells of a 24-well plates. Cells were fixed for 20 minutes with fixing solution (PBS with 3.7% (v/v) paraformaldehyde (Thermo Scientific, Cat #28906)). PFA was quenched and washed with TBS and the samples were then permeabilized and blocked for one hour with Imaging Buffer (PBS, 3% BSA, and 0.1% TritonX-100). Primary antibodies were added at 1:1000 in fresh Imaging Buffer. After labeling for 1 hour at room temperature, the coverslips were washed 3X with PBS and secondary antibodies were added at 1:1000 in Imaging Buffer. After labeling for ~45 minutes, the samples were washed 3X with PBS and cover slips were mounted on glass slides with Prolong Gold Mounting Media (Thermo Scientific, Cat #P36934) and allowed to dry overnight. Samples were imaged on a Zeiss LSM 900 with Airyscan 2. Following Airyscan processing in Zen Blue software (Zeiss), images were processed in FIJI.

### Fixed Imaging: Validation of RME-8 Antibody

HEK293-FLP cells and HEK293-FLP cells expressing UBC:DOR-APEX2 were siRNA treated according to protocol above. HEK293-FLP cells were treated with non-targeting control siRNA while HEK293-FLP cells expressing UBC:DOR-APEX2 were treated with DNAJC13 siRNA. 24 h following transfection, cells were lifted and replated onto poly-L-lysine coated coverslips in wells of a 24-well plates at equal confluence (~25%). Fixation, staining, imaging, and processing were performed as described, using Prolong Gold Mounting Media with DAPI (Thermo Scientific, Cat# P36962).

### Pearson’s Correlation Coefficient

HEK293-FLP cells expressing UBC:DOR-APEX2 were stained for DOR, VPS, and RME as stated above. Pearson’s Correlation Coefficient was calculated for RME8/VPS35 and RME8/DOR using 3 images per slide (technical replicates) with 3 slides per condition (biological replicates). Pearson’s Coefficients were calculated using Zen Blue software following a background subtraction step. Background subtraction values were obtained by establishing 3 frames not including cellular material and averaging these intensity values.

### Scoring Adjacency of DOR, RME-8, and VPS35

HEK293-FLP cells expressing UBC:DOR-APEX2 were stained for DOR, VPS35, and RME-8 as stated above. Super resolution Airyscan images were processed in 3-D in Zen Blue and the images were adjusted for brightness and contrast in FIJI. Slides were fixed in biological triplicate and images were acquired from the two conditions (VPS35/RME-8 and DOR/RME-8). For DOR/RME-8, 10 images were acquired at 63X magnification per biological replicate summating to 30 images processed. As VPS35/RME-8 condition contained more scorable structures per image, 3-5 images were acquired at 63X magnification per biological replicate summating to 13 images processed. Images were processed and intracellular structures—in which both channels were in close proximity—were identified. Subsequently, all images were then randomized with a numerical code and given to a blinded reviewer for scoring of “adjacency.” Adjacency: <50% of the RME-8 signal colocalizes with the signal from either VPS35 or DOR. If more 50% of RME-8 signal colocalized, the structure was defined as “O” for overlap. Scoring was performed for data obtained from three independent biological replicates with 3-5 images per replicate.

### Surface Rendering in Imaris

Super resolution Airyscan images were processed in 3-D in Zen Blue and the file types were converted using Imaris File Converter. The Surfaces tool in Imaris 9.0 was utilized to produce 3-dimmensional surfaces for the three relevant channels (FLAG (DOR), RME-8, and VPS35). Surface grain size was set to 0.100 *μ*m and surfaces were rendered as partially opaque to view overlap between the channels.

### Sample Size and Biological Replicates

Sample size was predetermined for all experiments with the exception of Figure 4F in which an increasing rate of DOR degradation in the control condition caused us, *ad hoc*, to add an additional replicate. Biological replicates refer to experiments performed on different days with independently cultured cells, with the exception of Figure 2D in which a two of the replicates were performed from independently cultured cells in the morning and afternoon of the same day. For example, a stable cell line is kept in culture for one month and one to two experiments were performed on different days per week across several weeks. All biological replicates (referred to as “n”) were listed in the figure legends. All experiments were n=3 with the following exceptions: n=4 for S2A, S2C, 4F; n=2 for S1C, S1D, S4D; n=1 for 2A and S4E.

### Technical Replicates

Technical replicates refer to biological samples processed in parallel at the same day and time. All data generated using the APEX2/AUR-lysate assay used two averaged technical replicates per biological replicate with the exception of Figures 2D, 3D, 4A, and 4D which had three averaged technical replicates per biological replicate and S2B which did not have technical replicates.

### Statistics

Figure 2D, 4D, 4F, and S4C were analyzed using a one-way ANOVA, alpha=0.05, with Dunnett’s multiple comparison correction. Figures 1D, 1F, and S2F were analyzed with a two-tailed paired t-test. Figure 4A was analyzed with a multiple t-test, assuming the same SD for each population, with an alpha=0.05 and Holm-Sidak correction.

---

**Figure S1:**
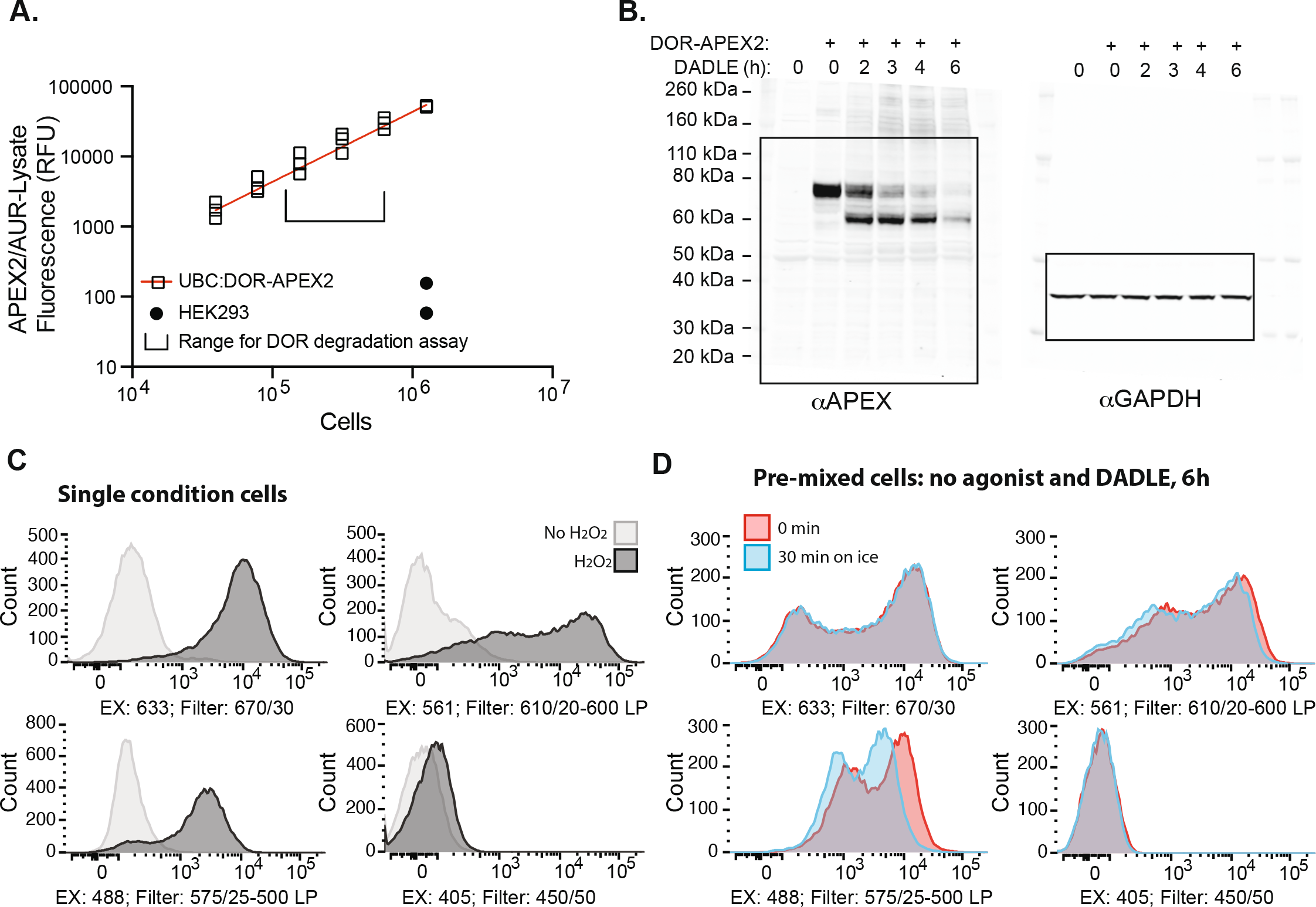
APEX2 as a fluorescence-based sensor for expression and lysosomal trafficking. **(a)** APEX2/AUR lysate reaction with specific numbers of HEK293-FLP cells stably expressing UBC:DOR-APEX2 or non-expressing control. Standard range for examining DOR trafficking to the lysosome for this cell line shown in brackets. N=3, linear regression fit to the data and constrained to the origin. **(b)** Uncropped western blot corresponding to Figure 1B. **(c)** Single cell analysis of APEX2/AUR intact assay (H_2_0_2_, dark grey) or no reaction conditions (no H_2_0_2_, light grey). Cells were analyzed on a BD FACS Aria II, n=2, representative example shown. **(d)** Single cell analysis of stable deposition of reaction product(s) of the APEX2/AUR assay in HEK293 cells stably expressing CMV:DOR-APEX2. Two populations of cells (unstimulated or stimulated for DADLE for 6 hours) were mixed prior to the APEX2/AUR assay, and analyzed on a BD FACS Aria II immediately following the reaction (red) or after 30 minutes on ice (blue). N=2, representative example shown.

**Figure S2:**
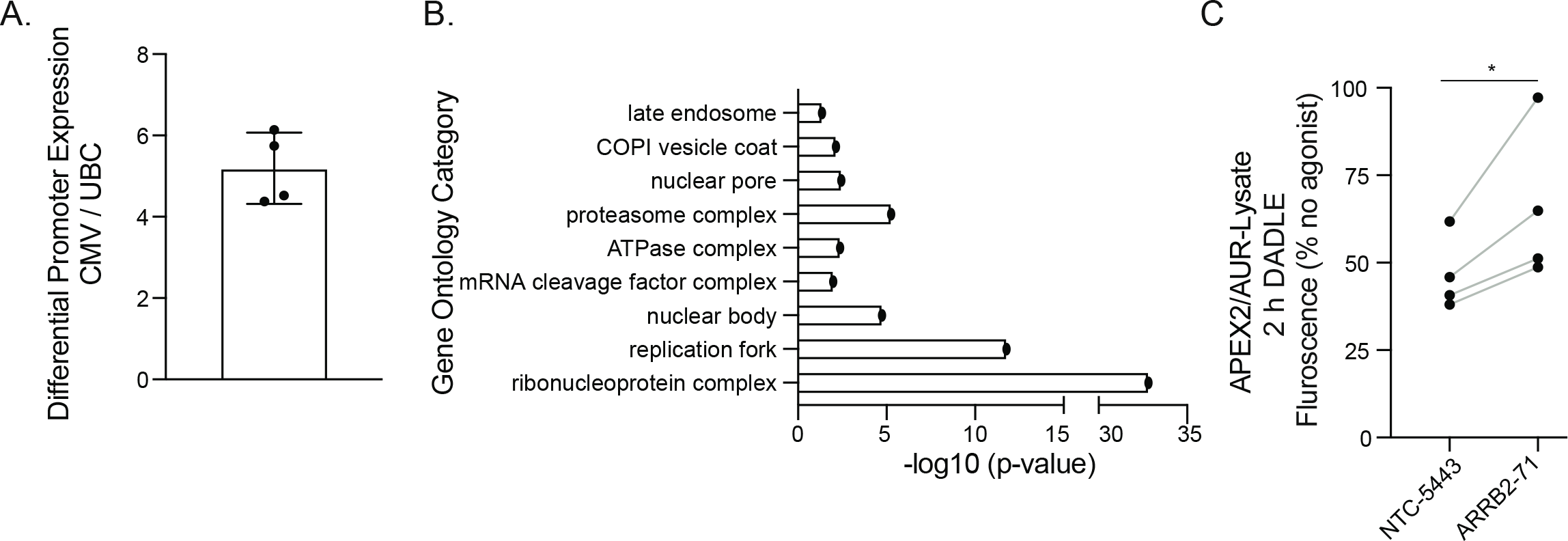
Genome-wide CRISPRi screen for genes affecting DOR expression and trafficking. **(a)** Relative surface expression of DOR-APEX2 stably expressed from the CMV or UBC promoter, probed for surface immunofluorescence with anti-FLAG M1 647, and analyzed with a Beckman Coulter Cytoflex S, n=4. **(b)** Gene ontology overrepresentation test from the 492 hits from the CRISPRi screen analyzed using PANTHER analysis for “Slim Cellular Component” and plotted based on calculated p-value. **(c)** Effects of beta-arrestin 2 targeting sgRNA on DOR trafficking to the lysosome were analyzed in HEK293-FLP cells stably expressing UBC:DOR-APEX2 and dCas9-BFP-KRAB, n=4, and data were analyzed using a two-tailed paired t-test (p=0.0485).

**Figure S3:**
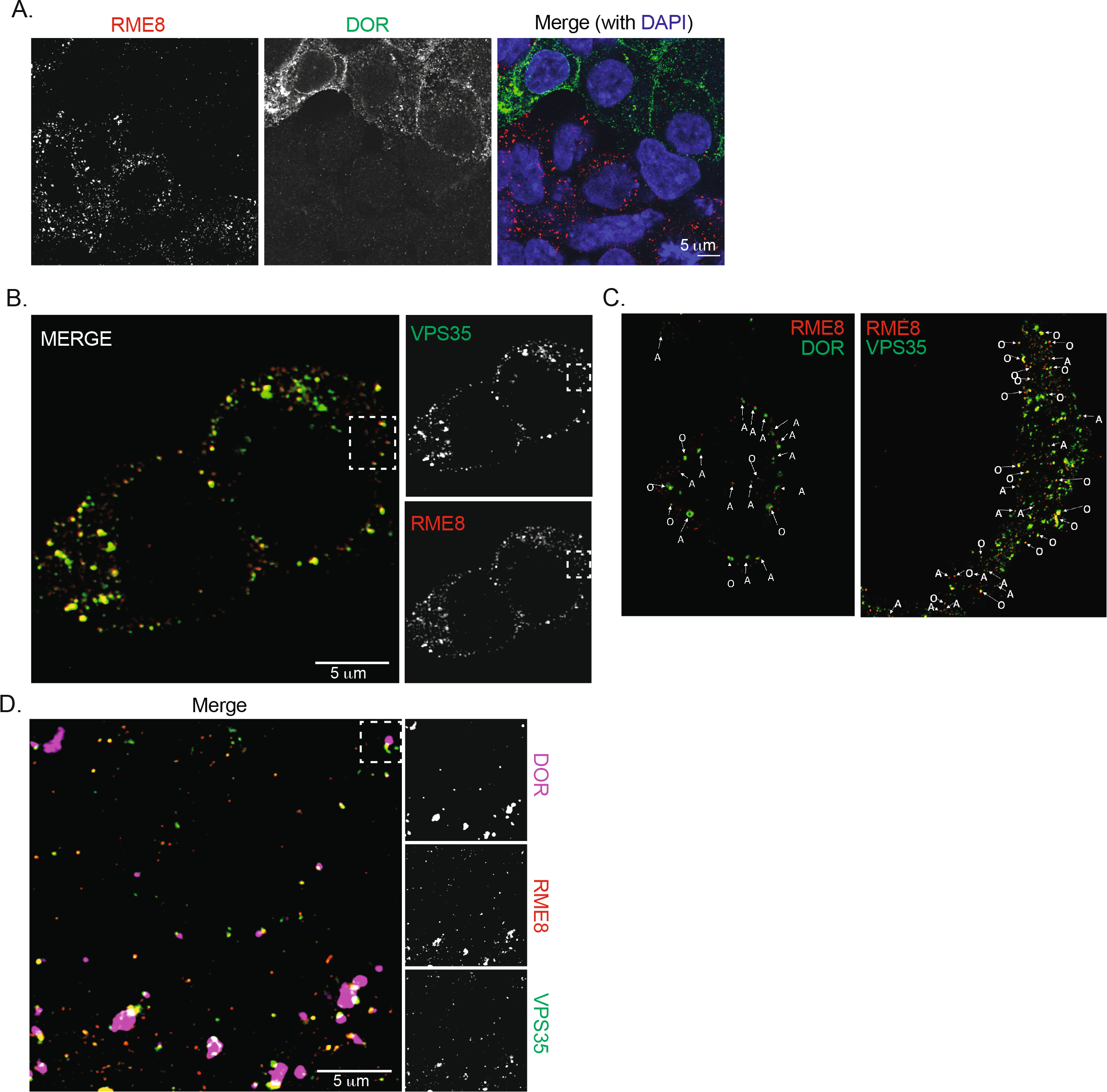
RME-8 localizes to DOR positive endosomes. **(a)** Validation of rabbit anti-RME-8 antibody using two populations of cells independently transfected with control siRNA (HEK293-FLP) or a siRNA pool targeting RME-8 (HEK293 FLP UBC:DOR-APEX2). Populations were mixed 48 h after transfection, fixed and permeabilized at 72 h, and then probed with anti-FLAG (DOR), anti-RME8, and DAPI. Scale bar is 5 *μ*m, n=3. **(b)** Fixed and permeabilized HEK293-FLP cells stably expressing UBC:DOR-APEX2 were probed with anti-RME-8 and anti-VPS35. Scale bar 5 *μ*m, n=3, white box corresponds to “Example 1” in 3C. **(c)** An example of the scoring system used to quantify “adjacency” between VPS35/RME-8 and DOR/RME-8. **(d)** Fixed and permeabilized HEK293-FLP cells stably expressing UBC:DOR-APEX2 were probed to detect endogenous RME-8 with anti-FLAG (DOR), anti-RME-8, and anti-VPS35. Scale bar 5 *μ*m. N=3, white box corresponds to “Example 1” in 3F.

**Figure S4:**
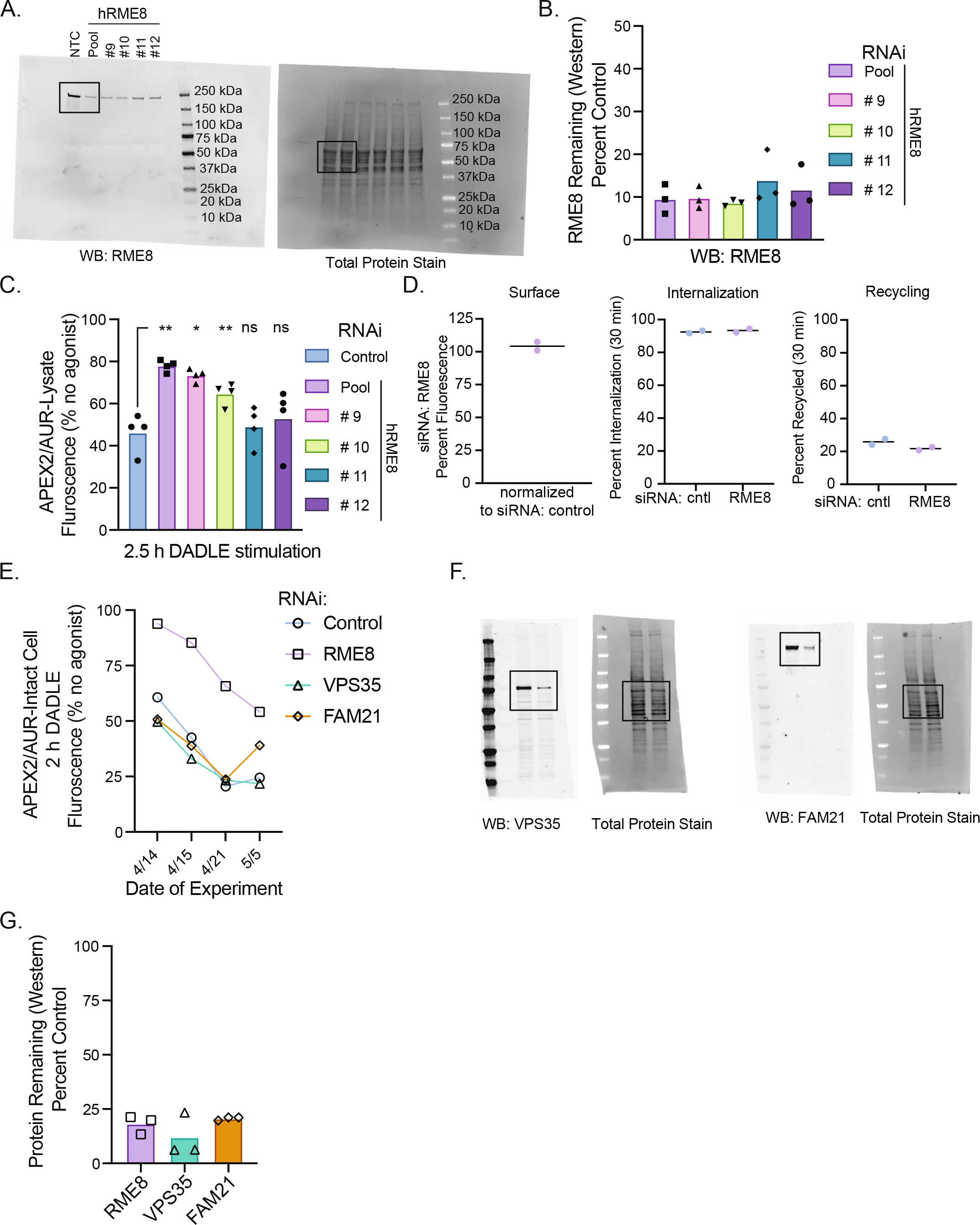
RME-8 regulates trafficking of DOR to the lysosome. **(a)** Uncropped western blot corresponding to Figure 4B. **(b)** Quantification of RME-8 knockdown as analyzed by western blot and mediated by a siRNA pool or its substituent individual siRNAs in HEK293 cells stably expressing UBC:DOR-APEX2. N=3 replicates were quantified using ImageLab (Bio-Rad) with the total protein stain used for sample loading normalization and each condition normalized to the NTC. **(c)** APEX2/AUR lysate assay following siRNA transfection with HEK293 cells stably expressing UBC:DOR-APEX2 with siRNA pool, or individual siRNAs, targeting RME-8 or a NTC. N=3 replicates were analyzed using a one-way ANOVA with Dunnett’s multiple comparisons test with an alpha of 0.05. Adjusted p-values vs control: pool (0.0091), 9 (0.0468), 10 (0.0076). **(d)** Surface expression of DOR at basal state or following internalization or recycling conditions was detected with anti-FLAG in HEK293-FLP cells stably expressing UBC:DOR-APEX2 and detected with a Beckman Coulter Cytoflex S, n=2. **(e)** Effects from knockdown of RME-8, FAM21, or VPS35 shown in 4F plotted here by date of experiment to demonstrate the change in degradation rate of DOR as a product of time in culture, n=1 per date. **(f)** Uncropped western blots corresponding to Figure 4C. **(g)** Quantification of RME-8, VPS35, or FAM21 knockdown as analyzed by western blot and mediated by gene-specific siRNA pools in HEK293 cells stably expressing UBC:DOR-APEX2. N=3 replicates were quantified using ImageJ with the total protein stain used for sample loading normalization and each condition normalized to the NTC.

## References

Becker, Martin, Heidi Noll-Puchta, Diana Amend, Florian Nolte, Christiane Fuchs, Irmela Jeremias, and Christian J. Braun. 2020. “CLUE: A Bioinformatic and Wet-Lab Pipeline for Multiplexed Cloning of Custom SgRNA Libraries.” Nucleic Acids Research 48 (13): e78.

Burke, M. D., and S. Orrenius. 1978. “The Effect of Albumin on the Metabolism of Ethoxyresorufin through O-Deethylation and Sulphate-Conjugation Using Isolated Rat Hepatocytes.” Biochemical Pharmacology 27 (11): 1533–38.

Cao, T. T., H. W. Deacon, D. Reczek, A. Bretscher, and M. von Zastrow. 1999. “A Kinase-Regulated PDZ-Domain Interaction Controls Endocytic Sorting of the Beta2-Adrenergic Receptor.” Nature 401 (6750): 286–90.

Chu Sin Chung, Paul, and Brigitte L. Kieffer. 2013. “Delta Opioid Receptors in Brain Function and Diseases.” Pharmacology & Therapeutics 140 (1): 112–20.

Civciristov, Srgjan, Cheng Huang, Bonan Liu, Elsa A. Marquez, Arisbel B. Gondin, Ralf B. Schittenhelm, Andrew M. Ellisdon, Meritxell Canals, and Michelle L. Halls. 2019. “Ligand-Dependent Spatiotemporal Signaling Profiles of the μ-Opioid Receptor Are Controlled by Distinct Protein-Interaction Networks.” The Journal of Biological Chemistry 294 (44): 16198–213.

Cullen, Peter J. 2008. “Endosomal Sorting and Signalling: An Emerging Role for Sorting Nexins.” Nature Reviews. Molecular Cell Biology 9 (7): 574–82.

Cullen, Peter J., and Florian Steinberg. 2018. “To Degrade or Not to Degrade: Mechanisms and Significance of Endocytic Recycling.” Nature Reviews. Molecular Cell Biology 19 (11): 679–96.

Damer, Cynthia K., Marina Bayeva, Emily S. Hahn, Javier Rivera, and Catherine I. Socec. 2005. “Copine A, a Calcium-Dependent Membrane-Binding Protein, Transiently Localizes to the Plasma Membrane and Intracellular Vacuoles in Dictyostelium.” BMC Cell Biology 6 (1): 46.

Degrandmaison, Jade, Khaled Abdallah, Véronique Blais, Samuel Génier, Marie-Pier Lalumière, Francis Bergeron, Catherine M. Cahill, et al. 2020. “In Vivo Mapping of a GPCR Interactome Using Knockin Mice.” Proceedings of the National Academy of Sciences of the United States of America 117 (23): 13105–16.

Dong, Chunmin, Catalin M. Filipeanu, Matthew T. Duvernay, and Guangyu Wu. 2007. “Regulation of G Protein-Coupled Receptor Export Trafficking.” Biochimica et Biophysica Acta 1768 (4): 853–70.

Duve, C. de, T. de Barsy, B. Poole, A. Trouet, P. Tulkens, and F. Van Hoof. 1974. “Commentary. Lysosomotropic Agents.” Biochemical Pharmacology 23 (18): 2495–2531.

Dwyer, Daniel J., Peter A. Belenky, Jason H. Yang, I. Cody MacDonald, Jeffrey D. Martell, Noriko Takahashi, Clement T. Y. Chan, et al. 2014. “Antibiotics Induce Redox-Related Physiological Alterations as Part of Their Lethality.” Proceedings of the National Academy of Sciences of the United States of America 111 (20): E2100–9.

Freeman, Caroline L., Geoffrey Hesketh, and Matthew N. J. Seaman. 2014. “RME-8 Coordinates the Activity of the WASH Complex with the Function of the Retromer SNX Dimer to Control Endosomal Tubulation.” Journal of Cell Science 127 (Pt 9): 2053–70.

Fujibayashi, Akemi, Tomohiko Taguchi, Ryo Misaki, Masashi Ohtani, Naoshi Dohmae, Koji Takio, Masashi Yamada, et al. 2008. “Human RME-8 Is Involved in Membrane Trafficking through Early Endosomes.” Cell Structure and Function 33 (1): 35–50.

Gendron, Louis, Catherine M. Cahill, Mark von Zastrow, Peter W. Schiller, and Graciela Pineyro. 2016. “Molecular Pharmacology of δ-Opioid Receptors.” Pharmacological Reviews 68 (3): 631–700.

Gilbert, Luke A., Max A. Horlbeck, Britt Adamson, Jacqueline E. Villalta, Yuwen Chen, Evan H. Whitehead, Carla Guimaraes, et al. 2014. “Genome-Scale CRISPR-Mediated Control of Gene Repression and Activation.” Cell 159 (3): 647–61.

Gilbert, Luke A., Matthew H. Larson, Leonardo Morsut, Zairan Liu, Gloria A. Brar, Sandra E. Torres, Noam Stern-Ginossar, et al. 2013. “CRISPR-Mediated Modular RNA-Guided Regulation of Transcription in Eukaryotes.” Cell 154 (2): 442–51.

Girard, Martine, and Peter S. McPherson. 2008. “RME-8 Regulates Trafficking of the Epidermal Growth Factor Receptor.” FEBS Letters 582 (6): 961–66.

Girard, Martine, Viviane Poupon, Francois Blondeau, and Peter S. McPherson. 2005. “The DnaJ-Domain Protein RME-8 Functions in Endosomal Trafficking.” The Journal of Biological Chemistry 280 (48): 40135–43.

Gomez-Lamarca, Maria, Laura Amy Snowdon, Ekatarina Seib, Thomas Klein, and Sarah Bray. 2015. “Rme-8 Depletion Perturbs Notch Recycling and Predisposes to Pathogenic Signaling.” The Journal of Cell Biology 210 (3): 517–517.

Guillen, Rodrigo X., Janel R. Beckley, Jun-Song Chen, and Kathleen L. Gould. 2020. “CRISPR-Mediated Gene Targeting of CK1δ/∊ Leads to Enhanced Understanding of Their Role in Endocytosis via Phosphoregulation of GAPVD1.” Scientific Reports 10 (1): 6797.

Guo, J., Y. Wu, W. Zhang, J. Zhao, L. A. Devi, G. Pei, and L. Ma. 2000. “Identification of G Protein-Coupled Receptor Kinase 2 Phosphorylation Sites Responsible for Agonist-Stimulated Delta-Opioid Receptor Phosphorylation.” Molecular Pharmacology 58 (5): 1050–56.

Han, Yisu, Tess Caroline Branon, Jeffrey D. Martell, Daniela Boassa, David Shechner, Mark H. Ellisman, and Alice Ting. 2019. “Directed Evolution of Split APEX2 Peroxidase.” ACS Chemical Biology 14 (4): 619–35.

Henry, Anastasia G., Ian J. White, Mark Marsh, Mark von Zastrow, and James N. Hislop. 2011. “The Role of Ubiquitination in Lysosomal Trafficking of δ-Opioid Receptors.” Traffic (Copenhagen, Denmark) 12 (2): 170–84.

Hermle, Tobias, Ronen Schneider, David Schapiro, Daniela A. Braun, Amelie T. van der Ven, Jillian K. Warejko, Ankana Daga, et al. 2018. “GAPVD1 and ANKFY1 Mutations Implicate RAB5 Regulation in Nephrotic Syndrome.” Journal of the American Society of Nephrology: JASN 29 (8): 2123–38.

Hislop, James N., Anastasia G. Henry, Adriano Marchese, and Mark von Zastrow. 2009. “Ubiquitination Regulates Proteolytic Processing of G Protein-Coupled Receptors after Their Sorting to Lysosomes.” The Journal of Biological Chemistry 284 (29): 19361–70.

Horlbeck, Max A., Luke A. Gilbert, Jacqueline E. Villalta, Britt Adamson, Ryan A. Pak, Yuwen Chen, Alexander P. Fields, et al. 2016. “Compact and Highly Active Next-Generation Libraries for CRISPR-Mediated Gene Repression and Activation.” ELife 5 (September). https://doi.org/10.7554/eLife.19760.

Johnson, Danielle E., Philip Ostrowski, Valentin Jaumouillé, and Sergio Grinstein. 2016. “The Position of Lysosomes within the Cell Determines Their Luminal PH.” The Journal of Cell Biology 212 (6): 677–92.

Kampmann, Martin. 2018. “CRISPRi and CRISPRa Screens in Mammalian Cells for Precision Biology and Medicine.” ACS Chemical Biology 13 (2): 406–16.

Kampmann, Martin, Michael C. Bassik, and Jonathan S. Weissman. 2013. “Integrated Platform for Genome-Wide Screening and Construction of High-Density Genetic Interaction Maps in Mammalian Cells.” Proceedings of the National Academy of Sciences of the United States of America 110 (25): E2317–26.

Kampmann, Martin, Michael C. Bassik, and Jonathan S. Weissman. 2014. “Functional Genomics Platform for Pooled Screening and Generation of Mammalian Genetic Interaction Maps.” Nature Protocols 9 (8): 1825–47.

Katayama, Hiroyuki, Akitsugu Yamamoto, Noboru Mizushima, Tamotsu Yoshimori, and Atsushi Miyawaki. 2008. “GFP-like Proteins Stably Accumulate in Lysosomes.” Cell Structure and Function 33 (1): 1–12.

Kouhen, O. M., G. Wang, J. Solberg, L. J. Erickson, P. Y. Law, and H. H. Loh. 2000. “Hierarchical Phosphorylation of Delta-Opioid Receptor Regulates Agonist-Induced Receptor Desensitization and Internalization.” The Journal of Biological Chemistry 275 (47): 36659–64.

Lacey, Joanne, Simon J. Webster, Paul R. Heath, Chris J. Hill, Lucinda Nicholson-Goult, Bart E. Wagner, Abdullah O. Khan, Neil V. Morgan, Michael Makris, and Martina E. Daly. 2022. “Sorting Nexin 24 Is Required for α-Granule Biogenesis and Cargo Delivery in Megakaryocytes.” Haematologica, January. https://doi.org/10.3324/haematol.2021.279636.

Lam, Stephanie S., Jeffrey D. Martell, Kimberli J. Kamer, Thomas J. Deerinck, Mark H. Ellisman, Vamsi K. Mootha, and Alice Y. Ting. 2015. “Directed Evolution of APEX2 for Electron Microscopy and Proximity Labeling.” Nature Methods 12 (1): 51–54.

Liu, Kai, Youli Jian, Xiaojuan Sun, Chengkui Yang, Zhiyang Gao, Zhili Zhang, Xuezhao Liu, et al. 2016. “Negative Regulation of Phosphatidylinositol 3-Phosphate Levels in Early-to-Late Endosome Conversion.” The Journal of Cell Biology 212 (2): 181–98.

Liu, Kai, Ruxiao Xing, Youli Jian, Zhiyang Gao, Xinli Ma, Xiaojuan Sun, Yang Li, et al. 2017. “WDR91 Is a Rab7 Effector Required for Neuronal Development.” The Journal of Cell Biology 216 (10): 3307–21.

Liu, Pei, Mikhail Khvotchev, Ying C. Li, Natali L. Chanaday, and Ege T. Kavalali. 2018. “Copine-6 Binds to SNAREs and Selectively Suppresses Spontaneous Neurotransmission.” The Journal of Neuroscience: The Official Journal of the Society for Neuroscience 38 (26): 5888–99.

Lobingier, Braden T., Ruth Hüttenhain, Kelsie Eichel, Kenneth B. Miller, Alice Y. Ting, Mark von Zastrow, and Nevan J. Krogan. 2017. “An Approach to Spatiotemporally Resolve Protein Interaction Networks in Living Cells.” Cell 169 (2): 350–360.e12.

Lobingier, Braden T., and Mark von Zastrow. 2019. “When Trafficking and Signaling Mix: How Subcellular Location Shapes G Protein-Coupled Receptor Activation of Heterotrimeric G Proteins.” Traffic 20 (2): 130–36.

Mann, Anika, Sophia Liebetrau, Marie Klima, Pooja Dasgupta, Dominique Massotte, and Stefan Schulz. 2020. “Agonist-Induced Phosphorylation Bar Code and Differential Post-Activation Signaling of the Delta Opioid Receptor Revealed by Phosphosite-Specific Antibodies.” Scientific Reports 10 (1): 8585.

Marchese, Adriano, May M. Paing, Brenda R. S. Temple, and Joann Trejo. 2008. “G Protein–Coupled Receptor Sorting to Endosomes and Lysosomes,” January. https://doi.org/10.1146/annurev.pharmtox.48.113006.094646.

Martell, Jeffrey D., Thomas J. Deerinck, Stephanie S. Lam, Mark H. Ellisman, and Alice Y. Ting. 2017. “Electron Microscopy Using the Genetically Encoded APEX2 Tag in Cultured Mammalian Cells.” Nature Protocols 12 (9): 1792–1816.

Martell, Jeffrey D., Thomas J. Deerinck, Yasemin Sancak, Thomas L. Poulos, Vamsi K. Mootha, Gina E. Sosinsky, Mark H. Ellisman, and Alice Y. Ting. 2012. “Engineered Ascorbate Peroxidase as a Genetically Encoded Reporter for Electron Microscopy.” Nature Biotechnology 30 (11): 1143–48.

Mitsui, Keiji, Yuri Koshimura, Yuriko Yoshikawa, Masafumi Matsushita, and Hiroshi Kanazawa. 2011. “The Endosomal Na(+)/H(+) Exchanger Contributes to Multivesicular Body Formation by Regulating the Recruitment of ESCRT-0 Vps27p to the Endosomal Membrane.” The Journal of Biological Chemistry 286 (43): 37625–38.

Nagy, Tamas, and Martin Kampmann. 2017. “CRISPulator: A Discrete Simulation Tool for Pooled Genetic Screens.” BMC Bioinformatics 18 (1). https://doi.org/10.1186/s12859-017-1759-9.

Norris, Anne, Prasad Tammineni, Simon Wang, Julianne Gerdes, Alexandra Murr, Kelvin Y. Kwan, Qian Cai, and Barth D. Grant. 2017. “SNX-1 and RME-8 Oppose the Assembly of HGRS-1/ESCRT-0 Degradative Microdomains on Endosomes.” Proceedings of the National Academy of Sciences of the United States of America 114 (3): E307–16.

Ostrowski, Philip P., Gregory D. Fairn, Sergio Grinstein, and Danielle E. Johnson. 2016. “Cresyl Violet: A Superior Fluorescent Lysosomal Marker.” Traffic (Copenhagen, Denmark) 17 (12): 1313–21.

Paek, Jaeho, Marian Kalocsay, Dean P. Staus, Laura Wingler, Roberta Pascolutti, Joao A. Paulo, Steven P. Gygi, and Andrew C. Kruse. 2017. “Multidimensional Tracking of GPCR Signaling via Peroxidase-Catalyzed Proximity Labeling.” Cell 169 (2): 338–349.e11.

Pfeiffer, Conrad T., Jialu Wang, Joao A. Paulo, Xue Jiang, Steven P. Gygi, and Howard A. Rockman. 2021. “Mapping Angiotensin II Type 1 Receptor-Biased Signaling Using Proximity Labeling and Proteomics Identifies Diverse Actions of Biased Agonists.” Journal of Proteome Research 20 (6): 3256–67.

Polacco, Benjamin J., Braden T. Lobingier, Emily E. Blythe, Nohely Abreu, Jiewei Xu, Qiongyu Li, Zun Zar Chi Naing, et al. 2022. “Profiling the Diversity of Agonist-Selective Effects on the Proximal Proteome Environment of G Protein-Coupled Receptors.” BioRxiv. https://doi.org/10.1101/2022.03.28.486115.

Popoff, Vincent, Gonzalo A. Mardones, Siau-Kun Bai, Valérie Chambon, Danièle Tenza, Patricia V. Burgos, Anbing Shi, et al. 2009. “Analysis of Articulation between Clathrin and Retromer in Retrograde Sorting on Early Endosomes.” Traffic (Copenhagen, Denmark) 10 (12): 1868–80.

Pradhan, Amynah A. A., Wendy Walwyn, Chihiro Nozaki, Dominique Filliol, Eric Erbs, Audrey Matifas, Christopher Evans, and Brigitte L. Kieffer. 2010. “Ligand-Directed Trafficking of the δ-Opioid Receptor in Vivo: Two Paths toward Analgesic Tolerance.” The Journal of Neuroscience: The Official Journal of the Society for Neuroscience 30 (49): 16459–68.

Puthenveedu, Manojkumar A., Benjamin Lauffer, Paul Temkin, Rachel Vistein, Peter Carlton, Kurt Thorn, Jack Taunton, Orion D. Weiner, Robert G. Parton, and Mark von Zastrow. 2010. “Sequence-Dependent Sorting of Recycling Proteins by Actin-Stabilized Endosomal Microdomains.” Cell 143 (5): 761–73.

Qi, Lei S., Matthew H. Larson, Luke A. Gilbert, Jennifer A. Doudna, Jonathan S. Weissman, Adam P. Arkin, and Wendell A. Lim. 2013. “Repurposing CRISPR as an RNA-Guided Platform for Sequence-Specific Control of Gene Expression.” Cell 152 (5): 1173–83.

Qin, Jane Yuxia, Li Zhang, Kayla L. Clift, Imge Hulur, Andy Peng Xiang, Bing-Zhong Ren, and Bruce T. Lahn. 2010. “Systematic Comparison of Constitutive Promoters and the Doxycycline-Inducible Promoter.” PloS One 5 (5): e10611.

Quirion, Béatrice, Francis Bergeron, Véronique Blais, and Louis Gendron. 2020. “The Delta-Opioid Receptor; A Target for the Treatment of Pain.” Frontiers in Molecular Neuroscience 13 (May): 52.

Rosenbluh, Joseph, Han Xu, William Harrington, Stanley Gill, Xiaoxing Wang, Francisca Vazquez, David E. Root, Aviad Tsherniak, and William C. Hahn. 2017. “Complementary Information Derived from CRISPR Cas9 Mediated Gene Deletion and Suppression.” Nature Communications 8 (1): 15403.

Ryder, Alan G., Sarah Power, and Thomas J. Glynn. 2003. “Fluorescence-Lifetime-Based PH Sensing Using Resorufin.” In Opto-Ireland 2002: Optics and Photonics Technologies and Applications, edited by Thomas J. Glynn. SPIE. https://doi.org/10.1117/12.463983.

Ryu, Seung W., Rose Stewart, D. Chase Pectol, Nicolette A. Ender, Oshadi Wimalarathne, Ji-Hoon Lee, Carlos P. Zanini, et al. 2020. “Proteome-Wide Identification of HSP70/HSC70 Chaperone Clients in Human Cells.” PLoS Biology 18 (7): e3000606.

Scherrer, Grégory, Petra Tryoen-Tóth, Dominique Filliol, Audrey Matifas, Delphine Laustriat, Yu Q. Cao, Allan I. Basbaum, et al. 2006. “Knockin Mice Expressing Fluorescent Delta-Opioid Receptors Uncover G Protein-Coupled Receptor Dynamics in Vivo.” Proceedings of the National Academy of Sciences of the United States of America 103 (25): 9691–96.

Semesta, Khairunnisa, Ruilin Tian, Martin Kampmann, Mark Zastrow, and Nikoleta Tsvetanova. 2021. “A High-throughput CRISPR Interference Screen for Dissecting Functional Regulators of GPCR/CAMP Signaling.” FASEB Journal: Official Publication of the Federation of American Societies for Experimental Biology 35 (S1). https://doi.org/10.1096/fasebj.2021.35.s1.02476.

Shen, Jinbo, Yonglun Zeng, Xiaohong Zhuang, Lei Sun, Xiaoqiang Yao, Peter Pimpl, and Liwen Jiang. 2013. “Organelle PH in the Arabidopsis Endomembrane System.” Molecular Plant 6 (5): 1419–37.

Shi, Anbing, Lin Sun, Riju Banerjee, Michael Tobin, Yinhua Zhang, and Barth D. Grant. 2009. “Regulation of Endosomal Clathrin and Retromer-Mediated Endosome to Golgi Retrograde Transport by the J-Domain Protein RME-8.” The EMBO Journal 28 (21): 3290–3302.

Sokolina, Kate, Saranya Kittanakom, Jamie Snider, Max Kotlyar, Pascal Maurice, Jorge Gandía, Abla Benleulmi-Chaachoua, et al. 2017. “Systematic Protein-Protein Interaction Mapping for Clinically Relevant Human GPCRs.” Molecular Systems Biology 13 (3): 918.

Tanowitz, Michael, and Mark von Zastrow. 2004. “Identification of Protein Interactions by Yeast Two-Hybrid Screening and Coimmunoprecipitation.” Methods in Molecular Biology (Clifton, N.J.) 259: 353–69.

Temkin, Paul, Ben Lauffer, Stefanie Jäger, Peter Cimermancic, Nevan J. Krogan, and Mark von Zastrow. 2011. “SNX27 Mediates Retromer Tubule Entry and Endosome-to-Plasma Membrane Trafficking of Signalling Receptors.” Nature Cell Biology 13 (6): 715–21.

Tian, Ruilin, Mariam A. Gachechiladze, Connor H. Ludwig, Matthew T. Laurie, Jason Y. Hong, Diane Nathaniel, Anika V. Prabhu, et al. 2019. “CRISPR Interference-Based Platform for Multimodal Genetic Screens in Human IPSC-Derived Neurons.” Neuron 104 (2): 239–255.e12.

Towne, Victoria, Mark Will, Brent Oswald, and Qinjian Zhao. 2004. “Complexities in Horseradish Peroxidase-Catalyzed Oxidation of Dihydroxyphenoxazine Derivatives: Appropriate Ranges for PH Values and Hydrogen Peroxide Concentrations in Quantitative Analysis.” Analytical Biochemistry 334 (2): 290–96.

Ven, Anne L. van de, Karen Adler-Storthz, and Rebecca Richards-Kortum. 2009. “Delivery of Optical Contrast Agents Using Triton-X100, Part 1: Reversible Permeabilization of Live Cells for Intracellular Labeling.” Journal of Biomedical Optics 14 (2): 021012.

Webb, Bradley A., Francesca M. Aloisio, Rabab A. Charafeddine, Jessica Cook, Torsten Wittmann, and Diane L. Barber. 2021. “PHLARE: A New Biosensor Reveals Decreased Lysosome PH in Cancer Cells.” Molecular Biology of the Cell 32 (2): 131–42.

Wittrup, K. D., and J. E. Bailey. 1988. “A Single-Cell Assay of Beta-Galactosidase Activity in Saccharomyces Cerevisiae.” Cytometry 9 (4): 394–404.

Xhabija, Besa, and Panayiotis O. Vacratsis. 2015. “Receptor-Mediated Endocytosis 8 Utilizes an N-Terminal Phosphoinositide-Binding Motif to Regulate Endosomal Clathrin Dynamics.” The Journal of Biological Chemistry 290 (35): 21676–89.

Xing, Ruxiao, Hejiang Zhou, Youli Jian, Lingling Li, Min Wang, Nan Liu, Qiuyuan Yin, Ziqi Liang, Weixiang Guo, and Chonglin Yang. 2021. “The Rab7 Effector WDR91 Promotes Autophagy-Lysosome Degradation in Neurons by Regulating Lysosome Fusion.” The Journal of Cell Biology 220 (8). https://doi.org/10.1083/jcb.202007061.

Yamashiro, D. J., and F. R. Maxfield. 1987. “Kinetics of Endosome Acidification in Mutant and Wild-Type Chinese Hamster Ovary Cells.” The Journal of Cell Biology 105 (6 Pt 1): 2713–21.

Yang, Dehua, Qingtong Zhou, Viktorija Labroska, Shanshan Qin, Sanaz Darbalaei, Yiran Wu, Elita Yuliantie, et al. 2021. “G Protein-Coupled Receptors: Structure- and Function-Based Drug Discovery.” Signal Transduction and Targeted Therapy 6 (1): 7.

Zastrow, Mark von, and Alexander Sorkin. 2021. “Mechanisms for Regulating and Organizing Receptor Signaling by Endocytosis.” Annual Review of Biochemistry 90 (1). https://doi.org/10.1146/annurev-biochem-081820-092427.

Zhang, Xiaoqing, Feifei Wang, Xiaoqing Chen, Yuejun Chen, and Lan Ma. 2008. “Post-Endocytic Fates of Delta-Opioid Receptor Are Regulated by GRK2-Mediated Receptor Phosphorylation and Distinct Beta-Arrestin Isoforms.” Journal of Neurochemistry 106 (2): 781–92.

Zhang, Y., B. Grant, and D. Hirsh. 2001. “RME-8, a Conserved J-Domain Protein, Is Required for Endocytosis in Caenorhabditis Elegans.” Molecular Biology of the Cell 12 (7): 2011–21.

Zhou, M., Z. Diwu, N. Panchuk-Voloshina, and R. P. Haugland. 1997. “A Stable Nonfluorescent Derivative of Resorufin for the Fluorometric Determination of Trace Hydrogen Peroxide: Applications in Detecting the Activity of Phagocyte NADPH Oxidase and Other Oxidases.” Analytical Biochemistry 253 (2): 162–68.

